# Oxaliplatin-induced cardiotoxicity in mice is connected to the changes in energy metabolism in the heart tissue

**DOI:** 10.1101/2023.05.24.542198

**Authors:** Junwei Du, Leland C. Sudlow, Kiana Shahverdi, Haiying Zhou, Megan Michie, Thomas H. Schindler, Joshua D. Mitchell, Shamim Mollah, Mikhail Y. Berezin

**Author notes:** Authors contributed equally (Junwei Du, Leland C. Sudlow). **Funding:** NCI/NIH R01CA208623, R21CA269099.

## Abstract

Oxaliplatin is a platinum-based alkylating chemotherapeutic agent used for cancer treatment. At high cumulative dosage, the negative effect of oxaliplatin on the heart becomes evident and is linked to a growing number of clinical reports. The aim of this study was to determine how chronic oxaliplatin treatment causes the changes in energy-related metabolic activity in the heart that leads to cardiotoxicity and heart damage in mice. C57BL/6 male mice were treated with a human equivalent dosage of intraperitoneal oxaliplatin (0 and 10 mg/kg) once a week for eight weeks. During the treatment, mice were followed for physiological parameters, ECG, histology and RNA sequencing of the heart. We identified that oxaliplatin induces strong changes in the heart and affects the heart’s energy-related metabolic profile. Histological post-mortem evaluation identified focal myocardial necrosis infiltrated with a small number of associated neutrophils. Accumulated doses of oxaliplatin led to significant changes in gene expression related to energy related metabolic pathways including fatty acid (FA) oxidation, amino acid metabolism, glycolysis, electron transport chain, and NAD synthesis pathway. At high accumulative doses of oxaliplatin, the heart shifts its metabolism from FAs to glycolysis and increases lactate production. It also leads to strong overexpression of genes in NAD synthesis pathways such as *Nmrk2*. Changes in gene expression associated with energy metabolic pathways can be used to develop diagnostic methods to detect oxaliplatin-induced cardiotoxicity early on as well as therapy to compensate for the energy deficit in the heart to prevent heart damage.

**Significance Statement:** This study uncovers the detrimental impact of chronic oxaliplatin treatment on heart metabolism in mice, linking high accumulative dosages to cardiotoxicity and heart damage. By identifying significant changes in gene expression related to energy metabolic pathways, the findings pave the way for the development of diagnostic methods to detect oxaliplatin-induced cardiotoxicity at an early stage. Furthermore, these insights may inform the creation of therapies that compensate for the energy deficit in the heart, ultimately preventing heart damage and improving patient outcomes in cancer treatment.

## INTRODUCTION

Published data on cardiotoxicities reported during chemotherapy reveal that cardiac damages are associated with oxidative stress, mitochondrial dysfunction, apoptosis, inflammation, and damage to the myocardium (1, 2). The changes in heart metabolic activity associated with chemotherapy are less studied, even though it might place the patient on a path to morbidity via heart disease. Clinically common metabolic changes reported during common first line of chemotherapy drugs include decreased energy production, changes in body composition, decreased muscle mass, altered macronutrient metabolism and changes in hormonal balance (2, 3). These changes can lead to weight loss, fatigue, appetite changes, and other symptoms including cardiac cachexia (4). Although many of these alterations are not permanent and might resolve as soon as chemotherapy is stopped, they may have long-term effects on heart function and overall health, depending on the severity of the metabolic changes (5).

Oxaliplatin is the first line of defense in colorectal cancers and is used against other malignancies, including gastric, pancreatic, and advanced hepatocellular carcinomas. Oxaliplatin’s efficacy is limited by its off-target toxicity leading to many acute and chronic side effects that include chemotherapy induced peripheral neuropathy (CIPN), anemia, nausea, vomiting, ototoxicity, renal toxicity, and hypersensitivity (6-9). One of the typical, but less studied side effects of oxaliplatin is its adverse effect on the heart. Patients treated with oxaliplatin often experience rapid breathing, chest pain, increase in heart rate, and an irregular heartbeat. Although emergency situations in oxaliplatin-treated patients are relatively rare compared to other drugs like anthracyclines (10, 11) or 5-fluorouacil (12) it is a rising concern given the increasing number of patients treated with oxaliplatin and in combination with other drugs. A growing number of cases related to severe coronary and cardiotoxicity of oxaliplatin alone (13, 14) or together with 5-fluoruracil or FOLFOX (15-18) have been reported.

Chemotherapy-induced alteration of heart metabolism occurs relatively quickly compared to other metabolic disorders: within several months, which corresponds to the duration of chemotherapy, rather than years, as in diabetes or other metabolic disorders. One of the hallmarks of heart disorders associated with diabetes and other metabolic disorders is the metabolic switch from fatty acid (FA) oxidation - the primary energy source of the heart - to glycolysis. We envisioned that oxaliplatin affects the heart in a similar way, by inhibiting enzymes involved in FA metabolism and activating the glycolytic pathway. Such a switch can lead to increased production of toxic byproducts such as lactate and a decrease in energy production (19).

Using ECG, post-mortem histology, and RNA-seq, we demonstrated direct damage to the heart caused by oxaliplatin in otherwise healthy mice. Genetic analysis identified a number of differentially expressed genes (DEGs) related to impaired energy-related metabolic processes and pathways. We observed that a strong suppression of FA metabolism, along with the suppression of amino acids and ketone body catabolism led to the activation of glycolysis potentially leading to the excess of toxic lactate. We also found a strong activation of the NAD+ synthesis pathway. These results provide key insights into the regulation of metabolic flux during oxaliplatin treatment and suggest potential diagnostic and therapeutic interventions.

## RESULTS

### Body mass is affected with the accumulative dosage of oxaliplatin

Over the course of the treatments, body masses of the control group showed weight gain consistent with mouse ageing (**Figure 1**). At the end of the oxaliplatin treatment an average 15% loss of gain was observed in the 10 mg/kg group. The exact mechanism behind oxaliplatin-induced weight loss is not fully understood. However, it may be due to several factors such as loss of activity, gastrointestinal toxicity, decreased food intake, malabsorption, and significant changes in metabolism.

**Figure 1.**
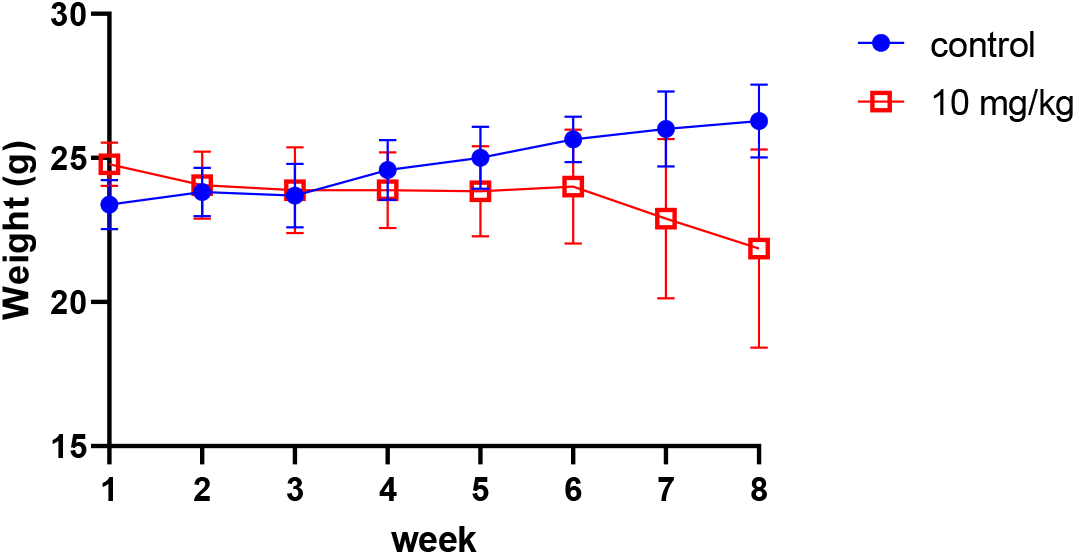
Weight loss of mice in the control and oxaliplatin treated mice. (Oxaliplatin, 10 mg/kg x 8 weekly injections, for control dextrose was used).

### High dose oxaliplatin decreases heart rates

Mouse heart rates were significantly affected at high-accumulated oxaliplatin dosage. Heart rates (HR) of the anesthetized, control mice for weeks 0-8 averaged 416 ± 45 bpm well within the range of published data for isoflurane anesthetized mice (20) (**Figure 2**). The average HR of the 10 mg/kg group at week 8 was 305.5 ± 74.6 bpm. In particular, one of the 10 mg/kg oxaliplatin treated mice at week 8 showed a heart rate of 198.6 ± 2.53 bpm.

**Figure 2.**
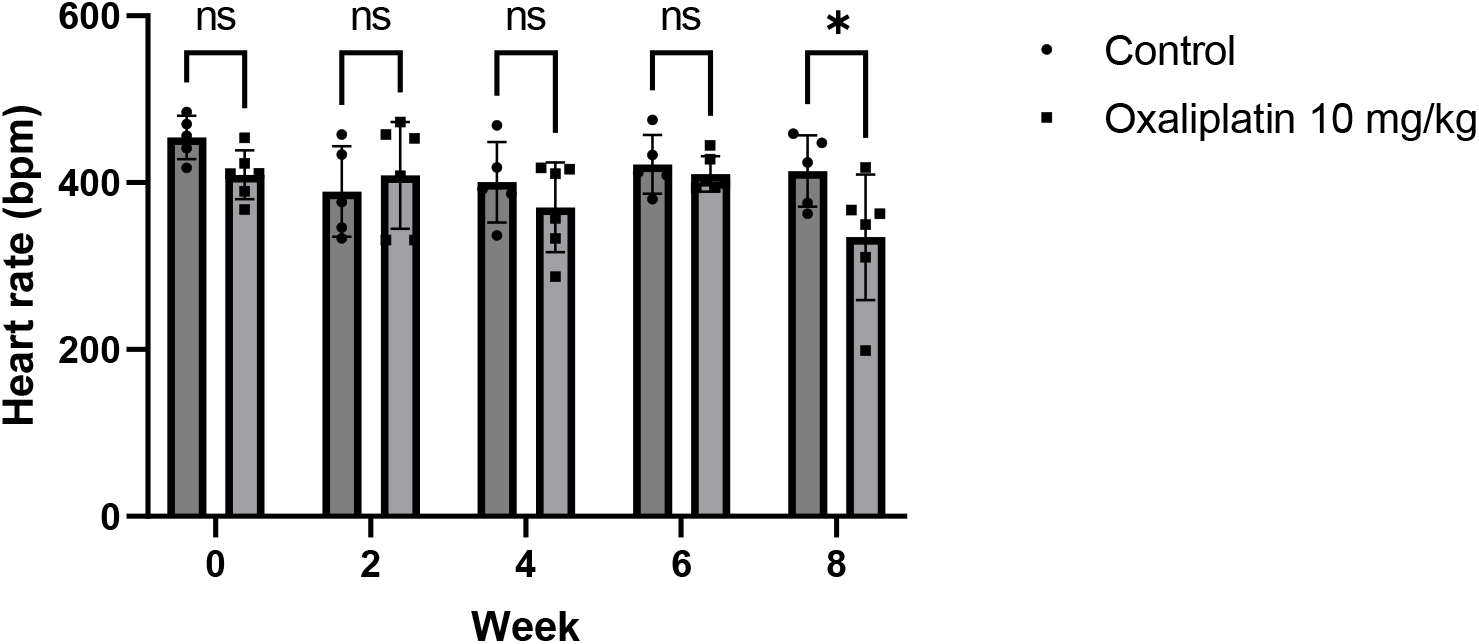
Effect of oxaliplatin on resting heart rates. By week 8, oxaliplatin-treated mice had a significantly lower resting heart rate. * p<0.05, Anova.

### Post-mortem histology reveals the damage to the heart even at low dosages

Varying degrees of heart damage in mice treated with oxaliplatin became evident during histological examination. Small areas of the heart showed inflammation associated with neutrophil and mononuclear cell infiltration (**Figure 3**). Regions of necrosis with loss of striations and necrotic coagulation and inflammation associated with the infiltration of a small number of neutrophils can also be seen.

**Figure 3.**
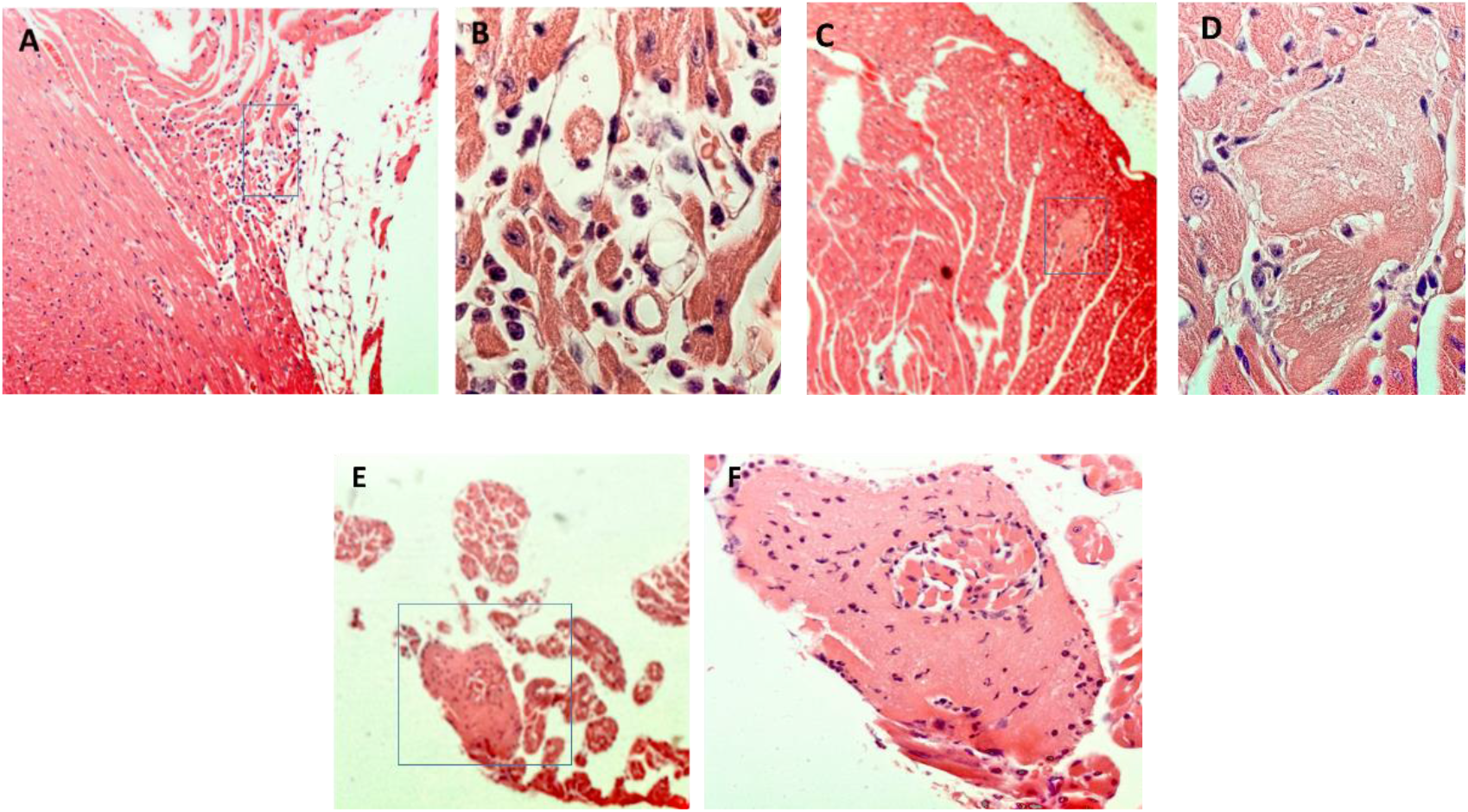
Histology of the heart of animal treated with oxaliplatin. **A:** Right atrial histology from a mouse treated with oxaliplatin for 8 weeks. Inset box is magnified in **B**. There is minimal interstitial infiltration by neutrophils and mononuclear cells; **C**: Ventricular histology near the apex of the mouse heart. Insert box is magnified in **D**. Focal myocardial necrosis is evident. This is a small focus of coagulative necrosis, with loss of cross-striations and loss of nuclei; **E:** Ventricular histology from near the right atrial appendage. Inset box is magnified in **F**. Focal myocardial necrosis is evident in the right atrial appendage that also shows infiltration of a small number of associated neutrophils.

### Oxaliplatin induces hypertrophy of the heart

The heart weight/body weight (HW/BW) index, whose normal value is around 0.4 – 0.45 mg/g for wild mice showed a dose dependent increase compared to the control mouse reaching 0.8 mg/g for the most affected mice (**Figure 4)**. The morphology of the heart in oxaliplatin mice resembled dilated cardiac hypertrophy that has been described in small animals in various cardiovascular diseases such as hypertension, myocardial infarction, valvular heart disease, heart fibrosis and hypertrophic cardiomyopathy (21, 22).

**Figure 4.**
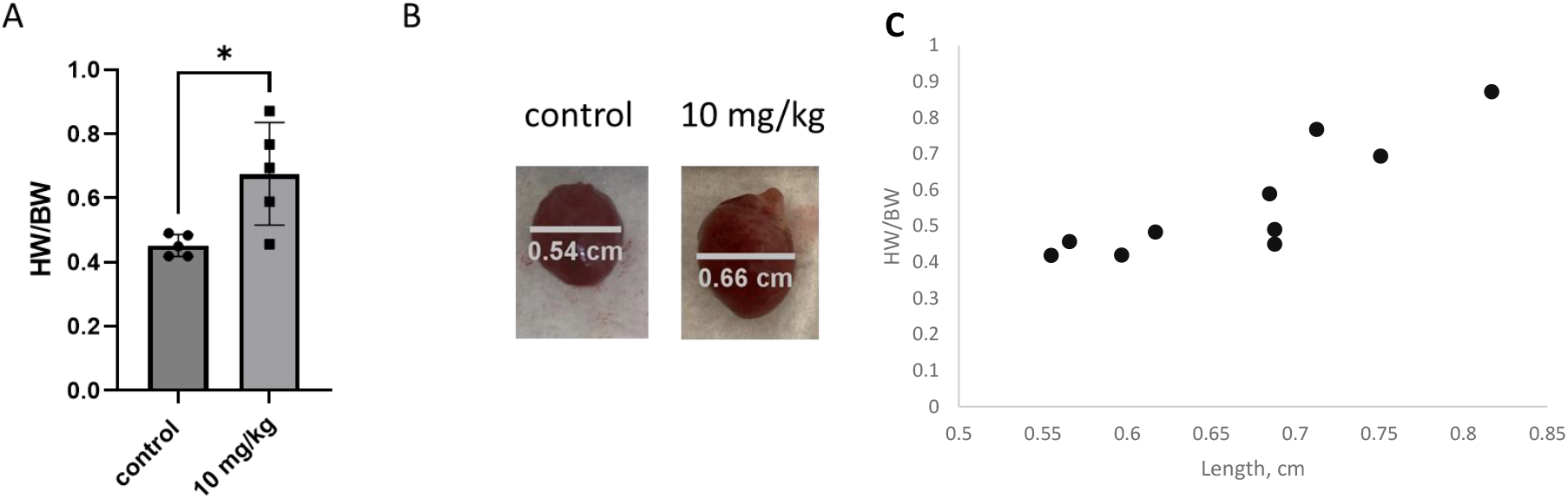
Gross effect of oxaliplatin on the heart. **A:** Heart weight/body weight (HW/BW, mg/g) index for mice treated with oxaliplatin. **B:** Examples images of the enlarged heart measured at the atrio-ventricular sulcus. **C:** Correlation between the heart size and the Heart weight/body weight index. The Pearson correlation between the size and the HW/BW = 0.86

### Markers of heart damage are overexpressed by qPCR

qPCR from the heart tissues showed elevated expression of several heart related disease markers including *Nppb* (b-type natriuretic peptide, a cardiac hypertrophic marker that is often used to diagnose and monitor heart failure), *Cdk1* (cyclin-dependent kinase 1, key cell cycle regulator), and *Lgals3* (galectin-3, whose expression is upregulated in inflammatory conditions and has been associated with a variety of cardiovascular diseases including heart failure and atherosclerosis) (**Figure 5**). Elevated levels of these genes indicate cardiac damage and inflammation and their expression is known to be upregulated in response to cardiotoxic agents and hemodynamic stress (23-26). In particular, *Lgals3* has been shown to be a useful marker for detecting cardiotoxicity in cancer patients undergoing treatment with certain chemotherapy drugs (24, 27). At the same time, cardiac troponin (Tnnt2), the conventional biomarker for myocardial infarction, was not associated with cardiotoxicity since *Tnnt2* differential expression was not statistically significant (**Figure S1**).

**Figure 5.**
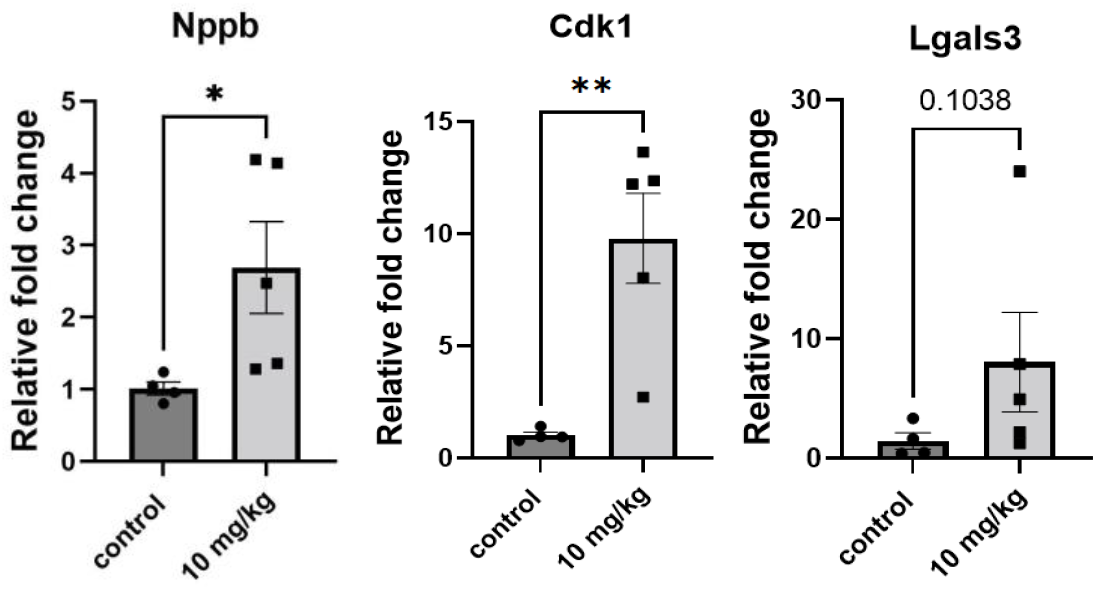
Changes in expression of markers of heart damage in mice by qPCR-RT in oxaliplatin (10 mg/kg, 8 dosages weekly) vs vehicle (control).

### RNA-seq demonstrates that heart metabolism is strongly affected by oxaliplatin

Following the establishment of oxaliplatin-induced cardiotoxicity, we investigated the differential expression of genes in the heart of control and oxaliplatin-treated mice using bulk RNA-seq of the entire heart. The RNA-seq data indicated that oxaliplatin induced substantial gene expression changes. From 13,775 identified genes in the heart, we found 2,667 genes or 19.3% were differentially expressed (FDR<0.05) **Figure 6A**). Heatmaps for the gene expression of all DEGs in the heart tissue combined with the hierarchical clustering analysis demonstrated an excellent separation between the oxaliplatin treated and control groups (**Figure 6B)**. From the GO Biological Processes analysis, we found significant enrichment of metabolic processes as shown in **Figure 6**

**Figure 6.**
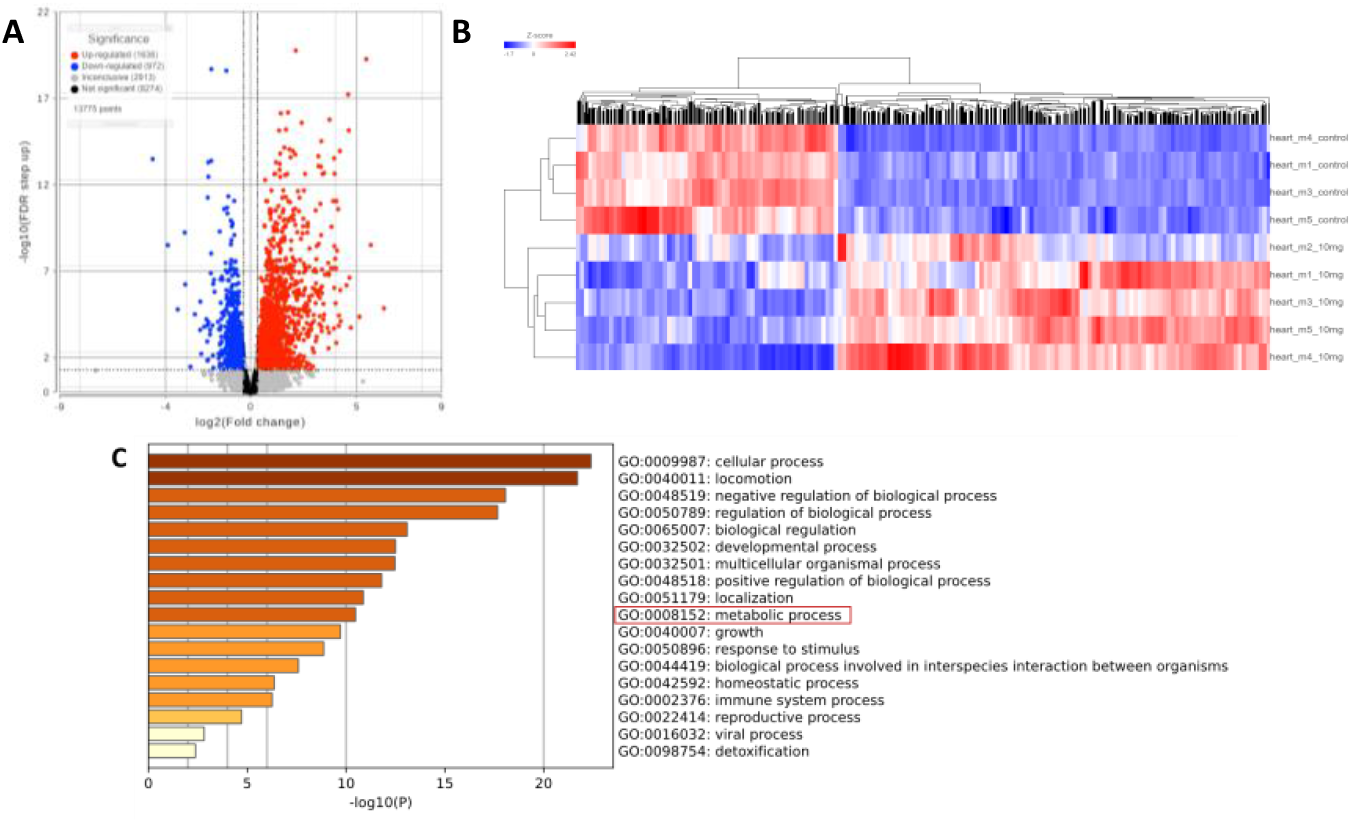
Effect of oxaliplatin on the RNA transcription in the heart. **A**: Volcano plot representation of genes from the heart tissue: Y threshold (FC) -1.25 to 1.25, X threshold FDR = 0.05. Total number of identified genes in the heart =13,755, upregulated =1,636, downregulated =972 (19.3% genes affected). **B**: Heatmap of all differentially expressed genes in the heart (total 2608): control dextrose-treated mice – top 4 rows, oxaliplatin -treated (bottom 5 rows). **C:** Top 20 processes affected by oxaliplatin by GO pathway enrichment analysis of all DEGs in the heart indicates metabolic pathway as one of the top affected pathways.

From the pool of affected metabolic genes, we narrowed our focus on energy metabolism. We have identified 75 genes directly involved in the nine major energy-related pathways including FA transport and oxidation, amino acid catabolism, ketone body catabolism, glycolysis, TCA cycle, oxidative phosphorylation, Cori cycle, NAD synthesis, and de novo synthesis of triglycerides. The full list of these pathways and genes is given in (Error! Reference source not found.). Genes that regulate metabolic genes were not included in the study.

Among the genes that broadly represent a variety of metabolic pathways (KEGG database identifies 1340 genes relevant to the entire metabolism (28)), 272 were differentially expressed. Using the pool of affected metabolic genes, we narrowed our focus on energy metabolism. We identified 75 genes directly involved in the major energy-related pathways. The top 20 of these processes related to the energy metabolic pathway are shown in **Figure 7A**. The details of energy metabolism pathways such as fatty acid degradation, glycolysis, amino acid catabolism, and NAD synthesis are provided below.

**Figure 7.**
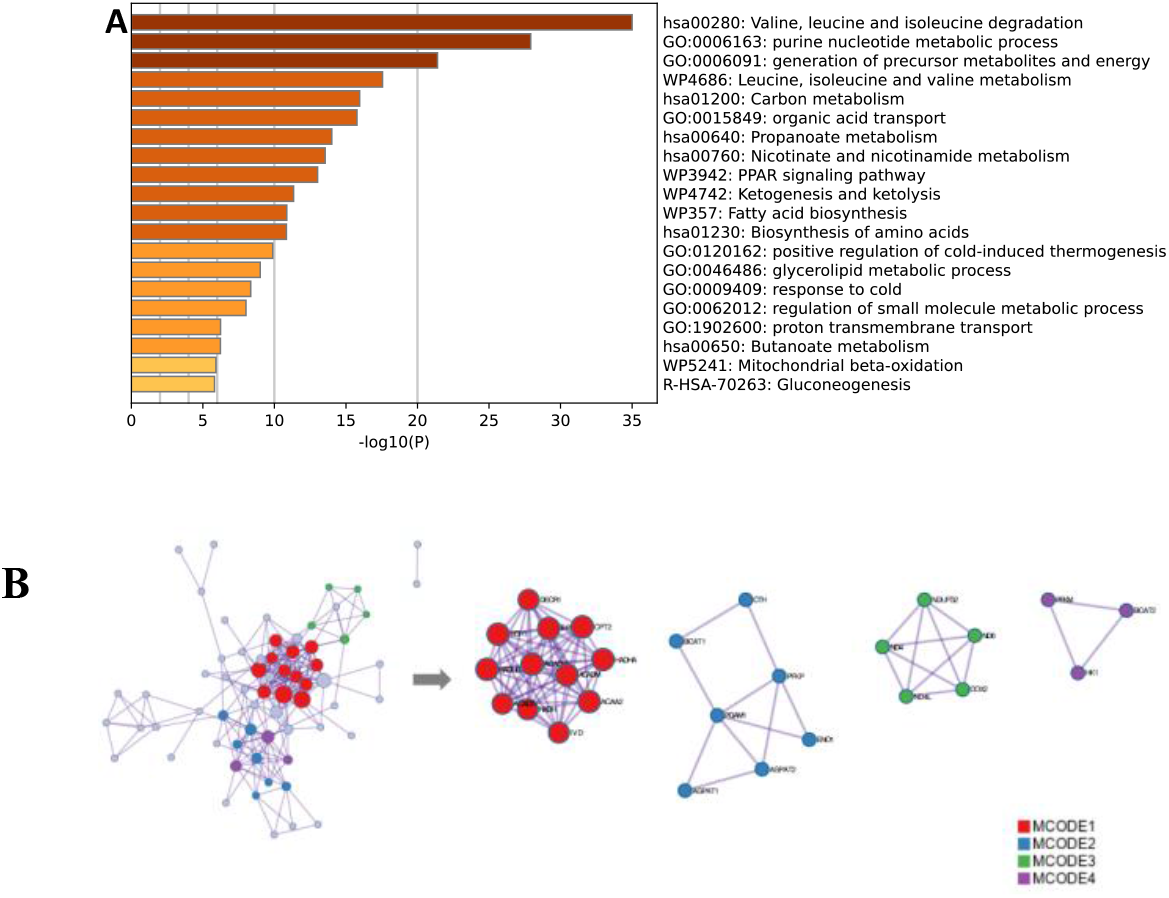
Effect of oxaliplatin on the RNA transcription of 75 metabolic DEGs the heart. **A:** Top 20 processes from the GO pathway enrichment analysis of 75 differentially expressed energy-related genes in the heart. **B:** Protein-protein interaction network and extracted 4 MCODE components/clusters. Network of enriched terms: (a) colored by cluster ID, where nodes that share the same cluster ID are typically close to each other; (b) colored by p-value, where terms containing more genes tend to have a more significant p-value. Cluster 1: *Acaa2, Acadm, Acads, Acadvl, Cpt2, Decr1, Eci1, Ech1, Hadh, Hadhb, Hadha, Ivd*; Cluster 2: *Adpat 1, Agpat 2, Bcat1,Cth, Eno1, Pgam1, Pfkp*; Cluster 3: *Ndufs2, Nd6, Nd4l, Nd4, Cox2*; Cluster 4: *Bcat2, Pfkm, Hk1*.

To further capture the relationships between the genes, a subset of enriched genes was selected and rendered as a network plot, where terms with a similarity >0.3 are connected by edges. The network was visualized with Cytoscape with “force-directed” layout and with edge bundled for clarity. In order to build a network of protein-protein interactions, only physical interactions in STRING (physical score > 0.132) and BioGrid were used. The resultant network contains a subset of proteins that form physical interactions with at least one other member in the list (**Figure 7B)**. The Molecular Complex Detection (MCODE) algorithm was then applied to this network to identify neighborhoods where proteins are densely connected (clusters). We identified four such clusters signifying four biological “functional modules”. Finally GO enrichment analysis was applied to each MCODE network to extract “biological meanings” from the network component, where the top three best p-value terms were retained (**Table 1**). The four functional modules represent the enrichment of the major energy-metabolisms pathways analyzed in detail below.

**Table 1.** Biological interpretation of the PPI network and MCODE components/clusters (functional modules).

| Network | Annotation | Value |
| --- | --- | --- |
| MyList | hsa00280: Valine, leucine and isoleucine degradation | 36.3 |
|  | GO:0032787: monocarboxylic acid metabolic process | 35.6 |
|  | GO:0044282: small molecule catabolic process | 34.3 |
| MCODE_ALL | GO:0044282: small molecule catabolic process | 25.9 |
|  | GO:0006635: fatty acid beta-oxidation | 25.7 |
|  | GO:0019395: fatty acid oxidation | 23.9 |
| MCODE_1 | GO:0006635: fatty acid beta-oxidation | 32.9 |
|  | GO:0019395: fatty acid oxidation | 31.2 |
|  | GO:0009062: fatty acid catabolic process | 30.9 |
| MCODE_2 | hsa01230: Biosynthesis of amino acids | 11.8 |
|  | GO:0006096: glycolytic process | 7.0 |
|  | GO:0006757: ATP generation from ADP | 7.0 |
| MCODE_3 | GO:0019646: aerobic electron transport chain | 12.7 |
|  | GO:0042775: mitochondrial ATP synthesis coupled electron transport: | 12.6 |
|  | GO:0042773: ATP synthesis coupled electron transport | 12.6 |
| MCODE_4 | GO:0044282: small molecule catabolic process | 5.8 |

### FA metabolic pathway is downregulated by oxaliplatin

Unlike other organs that rely heavily on glucose for energy, the healthy heart in humans and rodents relies predominantly on FAs. The healthy mouse heart utilizes 70% FAs and less than 30% carbohydrates as energy sources for ATP production (29). Small contributions also come from branched chain amino acid catabolism and ketone bodies. FAs utilization is primarily divided into the following steps: *i*) transport of FAs from capillary lumen by endothelium cells, *ii*) delivery of FAs from endothelium cells to cardiomyocyte membrane, *iii*) uptake of FAs into cardiomyocyte cytosol, *iv*) transport across the mitochondrial membrane with the following oxidation within the mitochondrial membrane. Many of the genes controlling these processes were downregulated in the treatment group (**Figure S2 and S3**).

#### i) FA transport from capillary to endothelium is upregulated

FAs transport from capillary to endothelium is the first step in delivery of FAs to cardiomyocytes, since the major source of free FAs or FAs bound to albumin is the blood plasma. FAs used by the heart come from the triglycerides (TGs) contained in TG-rich lipoproteins (chylomicrons and very low-density lipoproteins) or FAs bound to serum albumin in the blood (30) (**Figure 8A**). Highly abundant lipase LPL (encoded by *Lpl*) exported by cardiomyocytes hydrolyze the TGs from the lipoproteins (31, 32). The exported LPLs bind to glycosylphosphatidylinositol-anchored protein (GPIHBP1, encoded by *Gpihbp1*) on the luminal surface of the capillary endothelia. LPL binds chylomicrons and very-low density lipoproteins (VLDL) hydrolyzing FAs from TGs. While the level of LPL protein remains unchanged in the oxaliplatin-treated group, *Gpihbp1* was strongly overexpressed (FC=2.51), effectively increasing the concentration of LPL and therefore the concentration of FAs available for transport.

**Figure 8.**
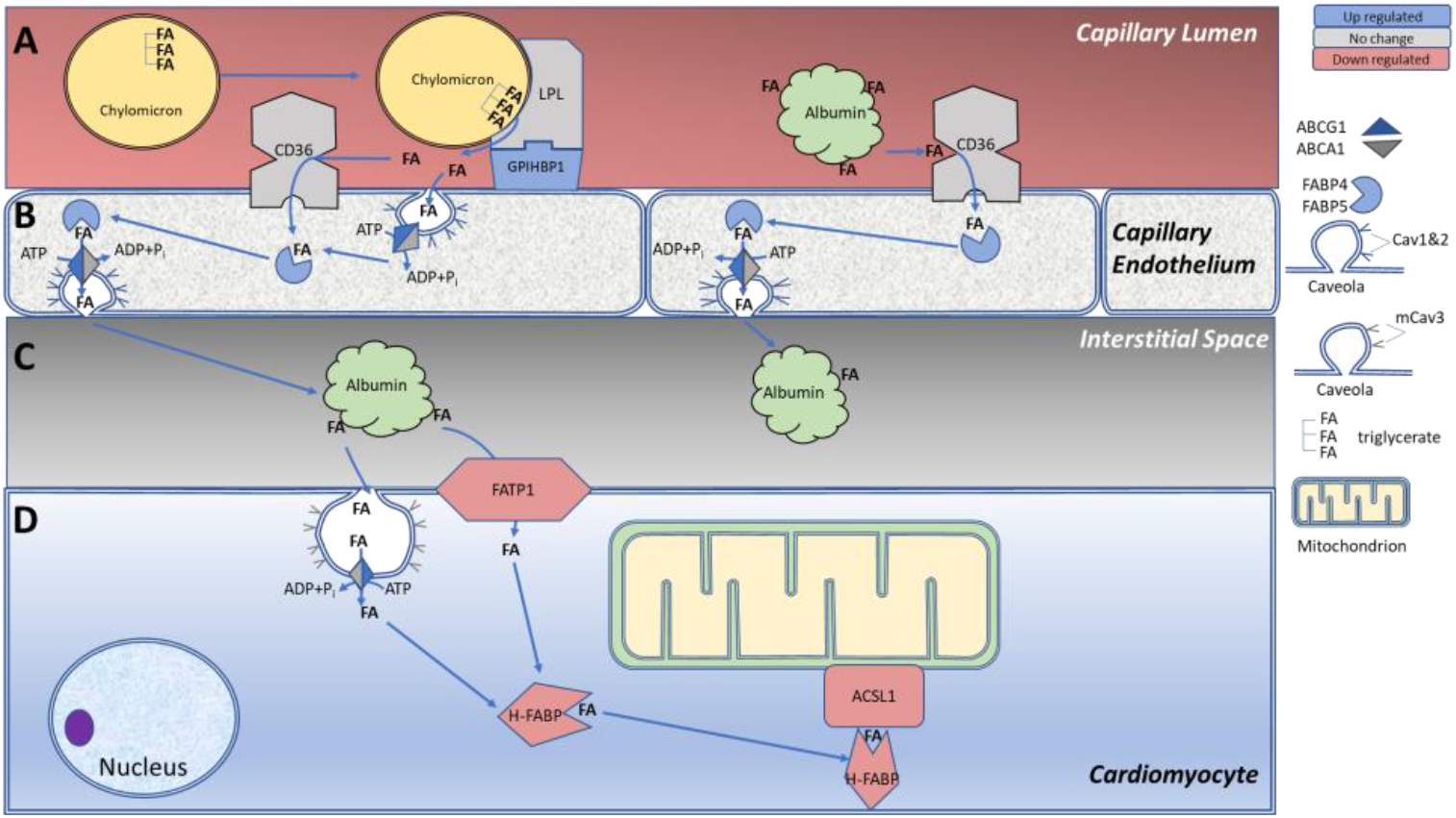
FA transport from capillary to the cytoplasm of cardiomyocytes affected by oxaliplatin. **A:** Lipoprotein lipase, LPL (*Lpl*) is transferred to the luminal side of the capillary endothelium where it binds to glycosylphosphatidylinositol-anchored protein 1 GPIHBP1 (*Gpihbp1)*. LPL hydrolyzes TGs contained in the chylomicrons, VLDLs producing free FAs. The released FAs are transported across the lumen-faced endothelium membrane by CD36 FA transporter (*Cd36*), or a flip-flop mechanism controlled by flippases ABCA1 and ABCG1 (*Abca1* and *Abcg1*). **B:** In the capillary endothelium, free FAs are picked up by FA binding proteins 4 and 5, FABP4,5 (*Fabp4* and *Fabp5*) and delivered to the interstitial space via a flip-flop mechanism. **C:** Interstitial albumin binds FAs and transports them to the cardiomyocyte sarcolemma via the FA translocase transport proteins such as FATP1 (*Slc27a1*), or flip-flop transport. **D:** In the cardiomyocytes, the heart FA binding protein 3, H-Fabp (*Fabp3*) pick the free FAs and transports them and their esters from the cell membrane to mitochondria for oxidation. Blue – upregulated genes, grey – unaffected genes, red downregulated genes.

To reach cardiomyocytes, liberated FAs must be transported across the capillary endothelium (**Figure 8B**). Two flippases, ABCA1 and ABCG1, encoded by *Abca1* and *Abcg1* move free FAs across the luminal and abluminal membranes of endothelial caveolae (33). Expression of *Abca1* was not impacted while *Abcg1* was significantly upregulated (FC=2.29). The expression of caveolar proteins caveolin1 and caveolin 2 (*Cav1* and *Cav2*) were upregulated. Another epithelial transporting pathway based on CD36 translocase (*Cd36*) was not affected by oxaliplatin (p>0.05). Once in the capillary endothelial cytoplasm, FAs are transported to abluminal epithelial membrane by FA binding proteins FABP4 and FABP5, encoded by *Fabp4* and *Fabp5* (34). Both genes were upregulated in the treatment group. The overexpression of these genes (**Table S1, Supplemental Information**) positively correlates with *Gpihbp1* expression (Pearson correlation coefficients *P*_(*Fabp4* vs. *Gpihbp1)*_*=0*.*944, P*_*(Fabp5* vs. *Gpihbp1)*_*=0*.*984*, **Figure S4**) suggesting highly coordinated hydrolysis of TGs and transport of liberated FAs through the capillary endothelial.

In the interstitial space the cardiomyocytes are picked up by interstitial albumin (35) as shown in **Figure 8C** and released into the cardiomyocytes through the transporter FATP1 (*Slc27a1*) and again using flippases ABCA1 and ABCG1 associated with caveolae in cardiomyocyte membranes that are regulated exclusively by caveolin 3 (*Cav3*, not affected) (36, 37). (**Figure 8D**). As the FA level in the interstitial space increases, it would be expected that the *Slc27a1* expression is upregulated. However, it was downregulated (FC=−1.53) suggesting the uptake of the FAs through the cardiomyocyte membrane is likely to be decreased and leaving the flip-flop mechanism to compensate for the supply of FAs through the upregulation of *Abcg1*. Once in the cardiomyocyte, FAs and their esters are carried to mitochondria for oxidation by a highly expressed heart FA binding protein H-FABP (*Fabp3*) (38). Lower *Fabp3* expression (FC=−1.44) (**Figure S2**) suggests impaired FAs transport to mitochondria.

The increased expressions of the genes involved in the transport of FAs to cardiomyocytes such as *Gpihbp1, Abcg1, Cav1* and *Cav2* along with the potentially lower transport of FAs (via *Fabp3*) to mitochondria indicate the likely accumulation of FAs in the interstitial space. This is in line with published data from *Fabp3*-KO mice that displayed lipid accumulation in the heart (39). The apparent reason for this accumulation is a survival mechanism to ensure the heart has a constant supply of energy. Indeed, elevated *Gpihbp1*expression has been previously observed in fasting mice (40). A decrease in *Slc27a1* expression (**Figure S2**) can also be tied to a decrease in membrane transport of FAs into the cardiomyocyte cytoplasm. There was a strong negative correlation (*P*=−0.83, **Table S1**) between *Slc27a1* that encodes FATP1 that controls the entrance of FAs to cardiomyocytes and *Slc25a20* that encodes carnitine acylcarnitine translocase (CACT) located on the inner mitochondrial membrane (IMM) (**Figure 9**) that is responsible for acyl-carnitines being transported into mitochondria. The strong negative correlation between these gene expressions indicates that two transport proteins play opposing roles in the regulation of FAs metabolism in the heart of oxaliplatin treated mice.

**Figure 9.**
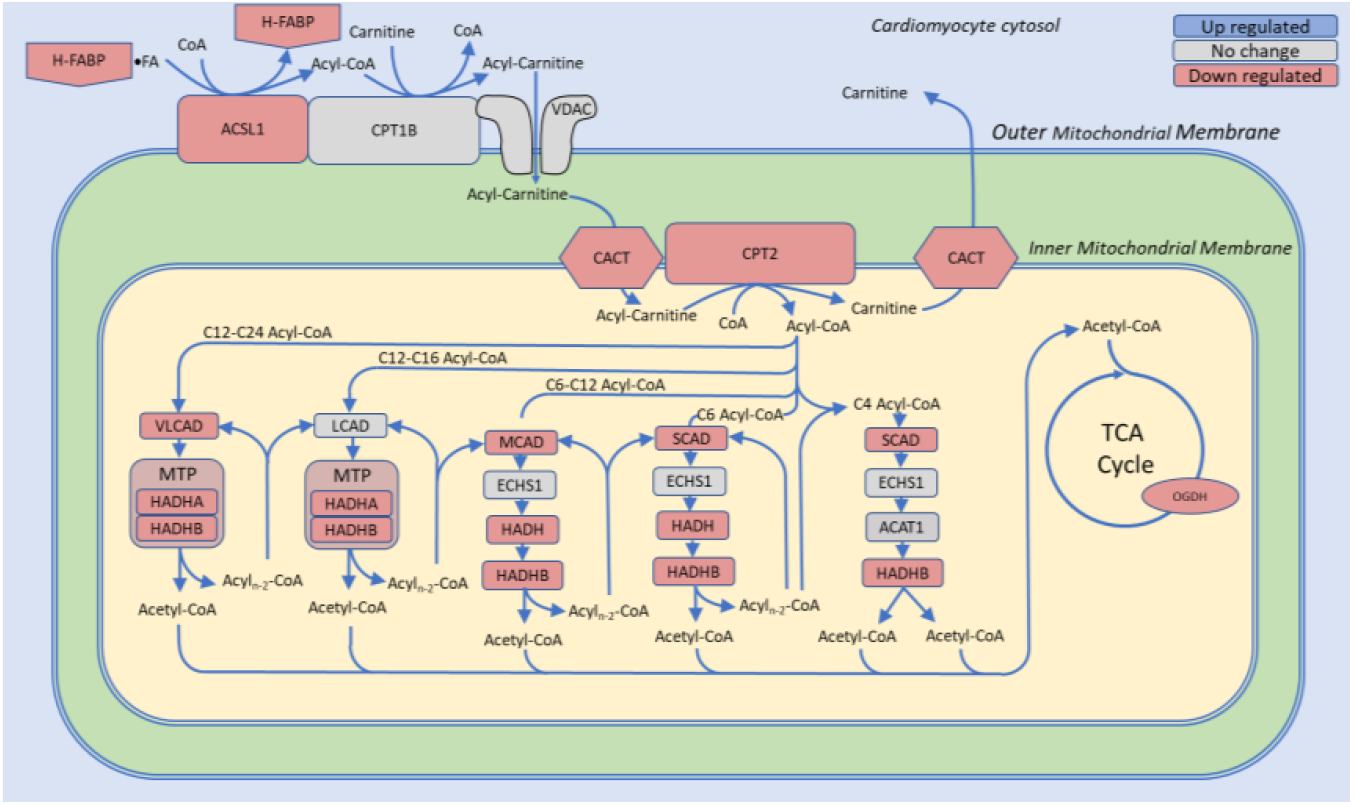
ß-oxidation pathway for fatty acids is strongly downregulated by oxaliplatin. Abbreviations: ACAT1, acetyl-CoA acetyltransferase 1 (*Acat1)*, ACSL1, acyl-CoA synthetase long chain family member 1 (*Acsl1*); CACT, Carnitine acylcarnitine translocase (*Slc25a20*); CPT1B, Carnitine O-palmitoyltransferase 1 (*Cpt1b*); CPT2, Carnitine O-palmitoyltransferase 2 (*Cpt2*); ECHS1, enoyl-CoA hydratase, short chain 1 (*Echs1*); HADH, hydroxyacyl-CoA dehydrogenase (*Hadh*); HADHA, hydroxyacyl-CoA dehydrogenase alpha and beta subunits of the mitochondrial trifunction protein complex (*Hadha* and *Hadhb*); H-FABP, heart fatty acid binding protein (*Fabp3*); LCAD, long chain acyl-CoA dehydrogenases (*Acadl*); MCAD, medium chain acyl-CoA dehydrogenase (*Acadm*); MTP, mitochondrial trifunction protein complex; SCAD, small chain acyl-CoA dehydrogenase (*Acads*); VLCAD, very long chain acyl-CoA dehydrogenases (*Acadvl*); VDAC, voltage dependent ion channels (*Vdac1, Vdac2*, or *Vdac3*). The pathway is derived from the KEGG database. Blue – upregulated genes, red downregulated genes, grey – not affected genes.

#### ii) FA transport through mitochondria membrane and β-oxidation pathways are downregulated

Following FAs import into the cardiomyocytes, FAs−H-FABP aggregates are transported to the outer mitochondrial membrane (OMM) (**Figure 9**). Because FAs cannot penetrate the OMM, FAs must first undergo conversion into acyl-CoA (41). This reaction is carried out by OMM-bound long chain synthetase ACSL1 encoded by *Acsl1* (42, 43) along with several other synthetases for different FA chain lengths. Downregulation of *Acsl1* (FC=−1.3) may suggest a lower conversion rate and less availability of the substrate to enter into mitochondria. A high correlation between the expression of *Fabp3* and *Acsl1* (*P*=0.951) (**Figure S3**) suggests tight coordination between the transport of the FA-derivative from the cytosol toward the mitochondria membrane and its conversion into acyl-CoA.

The formed acyl-CoA enters the acyl-carnitine pathway for transport of the FA across the OMM by first converting to acyl-carnitine by a transferase CPT1B and diffusion through a voltage dependent anion channel (VDAC) into the mitochondrial intermembrane space. From here, acyl-carnitines are transported through the inner mitochondrial membrane (IMM) by downregulated translocase CACT (encoded by *Slc25a20*, FC=−1.43) (44, 45) that exchanges cytoplasmic acyl-carnitines and mitochondrial free carnitines across the IMM (46). Lower levels of *Slc25a20* expression (**Figure S4**) may reflect the reduced level of substrates (FAs) in the cytosol and may lead to accumulation of the long-chain acyl-carnitines in the cytoplasm. This accumulation presents a serious condition that in extreme cases can lead to pathological changes in the heart causing arrhythmia and cardiac arrest (47).

Once the acyl-carnitines reach the mitochondrial matrix, the FA chain is re-transferred to CoA by a transferase CPT2 (*Cpt2*) (45) which is paired with CACT on the IMM (48). As a result of this process, acyl-CoA is formed again, and carnitine is returned to the cytoplasm via CACT. Predictably, *Cpt2* was downregulated (FC =−1.7). Given that this enzyme controls a rate limiting step in the β-oxidation pathway (49), the entire process of downstream β-oxidation would likely be substantially reduced. A high level of correlation between *Slc25a20* and *Cpt2* expressions (*P*=0.87) suggests that both processes (transport of acyl-carnitine and re-synthesis of acyl-CoA) are tightly coordinated.

β-oxidation of FAs into acetyl-CoA fragments and ATP is the final part of FA metabolism. This process is significantly affected by oxaliplatin with many enzymes and proteins being downregulated as illustrated in **Figure 9**. Both saturated FA and non-saturated FA β-oxidation pathways were affected. For saturated FAs, three out of four dehydrogenase enzymes that perform the first step to produce a corresponding enoyl-CoA were downregulated. That includes the variable chain dehydrogenases SCAD (*Acads*, FC=−1.47), MCAD (*Acadm*, FC=−1.43), and VLCAD (*Acadvl*, FC=−1.31) (**Figure S5**). Other downregulated enzymes on the saturated FA β-oxidation pathway include both subunits of the mitochondrial trifunction protein (MTP) made of dehydrogenases HADHA (*Hadha*, FC=−1.53), HADHB (*Hadhb*, FC*=*−1.47) and HADH (*Hadh*, FC= −1.5). These subunits perform multiple steps in β-oxidation and are essential for the normal functioning of the heart (50, 51). Due to the inability of these hydrogenases to catalyze the first step of the β-oxidation cycle of unsaturated FAs a different set of enzymes is employed (**Table S1)** (52). Similar to the saturated pathway, many of these genes (i.e., *Eci1, Decr1, Ech1*, (**Figure S6**) became downregulated due to oxaliplatin treatment.

Massive downregulation of multiple enzymes in the β-oxidation pathway along with the upregulation of FA transport from the capillaries to the interstitial space suggests an accumulation of FAs and their derivatives within cells, which can be toxic and contribute to heart failure (53). Failing to process FAs in sufficient quantities, the heart would be deprived of a significant source of energy to synthesize ATP. Since cardiomyocytes are unable to take up adequate levels of FAs, the heart must utilize other sources of energy such as amino acid catabolism, ketone body catabolism or switch to glycolysis as described in the following sections.

### Amino acid catabolism is downregulated

Amino acid catabolism represents a small source (less than 2%) of metabolic energy that comes from deaminated amino acids that enter the TCA cycle. Much of the output of the amino acid catabolism pathway is derived from branched chain amino acids (BCAAs) such as leucine, isoleucine, and valine (**Figure S7**) and neutral amino acids (NAA) glutamine, alanine, serine, glycine, cysteine and methionine (**Figure S8** – **S9**). The final products of these catabolic processes (acetyl-CoA, pyruvates and succinyl-CoA) enter the TCA cycle contributing to ATP generation.

Both BCAA and NAA pathways in the heart were mostly negatively affected by oxaliplatin. In the BCAA catabolic pathway, the transferase BCAT2 (*Bcat2*), which moves branched acids leucine, isoleucine into the mitochondria matrix and convert them to the keto-forms was downregulated (FC=−1.34) (54) (**Table S1, Figure S10**). Normally, keto products go through the dehydrogenase complex BCKDH, which has four subunits, in which both BCKDHA (*Bckdha*) and DBT (*Dbt*) were downregulated (both showed FC= −1.76). The genes that provide instruction for further dehydrogenation reactions performed by IVD (*Ivd*), MCAD (*Acadm*) and SCAD (*Acads*) were also downregulated with FC=−1.46, −1.44 and −1.47 correspondingly. Other downregulated genes in branched chain AA metabolism are shown in **Table S1**. The downregulation of these genes in the heart could potentially lead to impaired energy production due to decreased BCAA metabolism. Similar downregulation was also observed in the NAA catabolic pathways (**Figure S8** – **S9**). Several NAAs (i.e., alanine, glutamine, serine and some others) are capable of being converted into metabolic fuel when there is an insufficient supply of glucose or fatty acids in the body. Multiple enzymes involved in these processes were downregulated by oxaliplatin treatment. Overall, with both pathways negatively impacted by oxaliplatin, fewer acetyl-CoAs and other substrates via AA catabolism would be available for the TCA cycle.

### Ketone body metabolic routes are downregulated

Under stress, ketone bodies (acetone, acetoacetate, and β-hydroxybutyrate) can become primary energy sources for ATP production in the heart and as such a ketogenic diet is widely used to improve cardiac function (55). Ketone bodies are produced in the liver and transported across the myocardial cell membranes via the MCT1 channel (unaffected). The ketone body catabolism starts from transferring the CoA group from a donor succinyl-CoA molecule to the ketone body. This reaction is performed by a downregulated transferase OXCT1 (*Oxct1*, FC= −1.36) (**Figure S11**) that controls the rate limiting step of the entire ketone body catabolic process. The products are further processed by a transferase ACAT1 (*Acat1*, unaffected) into a pair of acetyl-CoA molecules. The resulting products are then processed by the TCA cycle and the Electron Transport Chain (ETC). Downregulation of *Oxct1* would restrict the heart’s ability to utilize ketone bodies resulting in decreased energy production and ketone bodies accumulation. An increase in ketones may lead to clinically dangerous ketoacidosis in patients with loss of appetite and lethargy (56).

### Tricarboxylic acid cycle (TCA) is weakly downregulated

Most metabolic energy pathways lead to the TCA cycle, which provides energy to cardiomyocytes. The TCA pathway relies on uninterrupted supply of pyruvate, NAD+ and FAD (**Figure 10**). TCA cycle genes were less affected by oxaliplatin. One noticeable exception was a downregulation of *Ogdh* (FC=−1.26) (**Figure S12**). The gene encodes the 2-oxoglutarate dehydrogenase (ODH) complex that catalyzes a rate-limiting step in the TCA cycle (57). The effect of this downregulation, although moderate, would slow the cycle, and potentially the entire metabolism.

**Figure 10.**
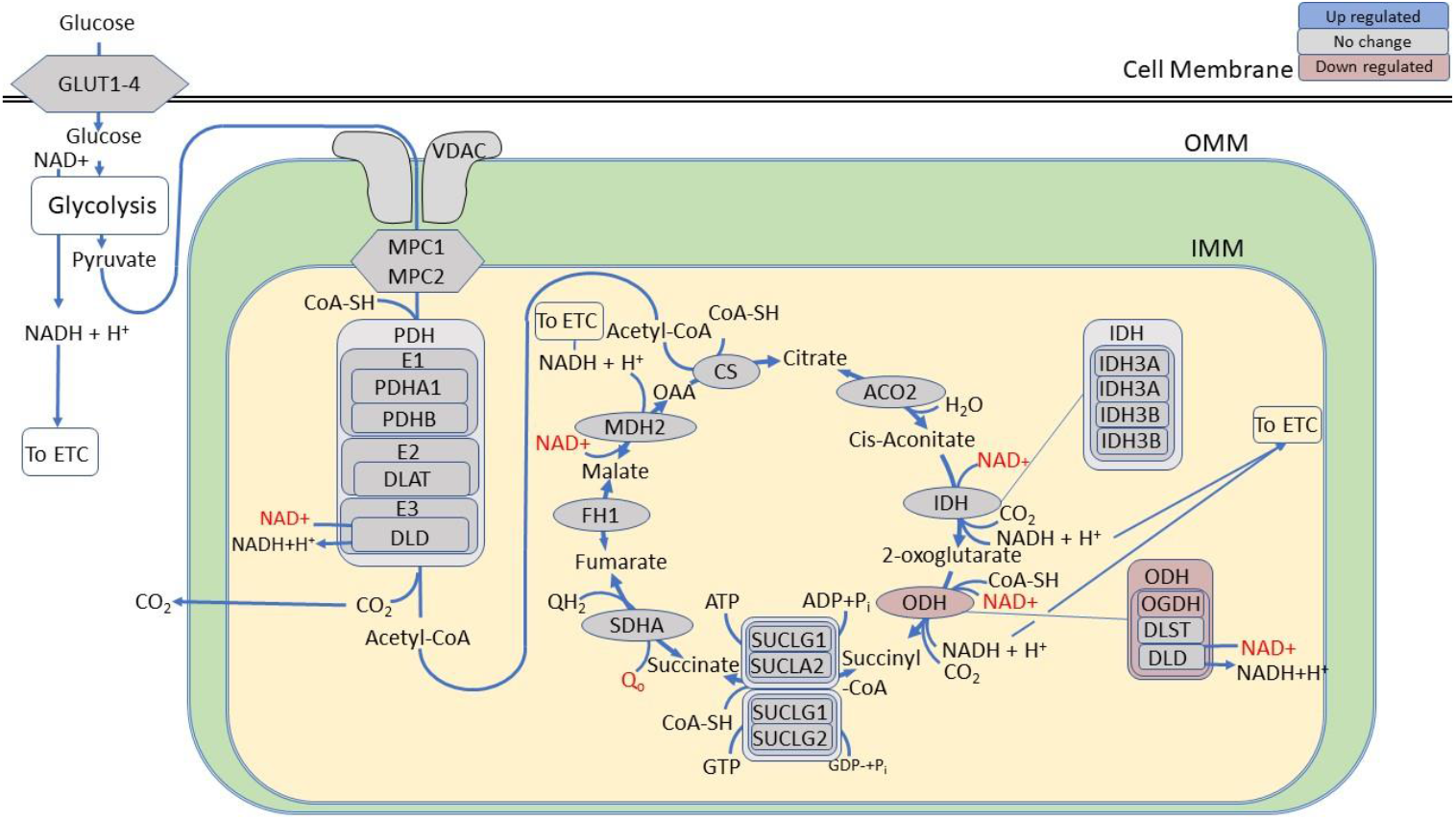
TCA cycle is affected by oxaliplatin. E1 component of 2-oxoglutarate dehydrogenase complex (OGDH) enzyme (*Ogdh)* exhibited statistically significant downregulation. The remaining two subunits of this complex encoded by *Dlst* and *Dld* and other genes controlling the TCA cycle were not affected. The other enzymes of the TCA cycle remained unaffected by oxaliplatin. That includes citrate synthase (*Cs*) that converts acetyl-CoA from β-oxidation of FAs or from glycolysis to form citrate, aconitase isomerases (*Aco1* and *Aco2*) that convert citrate to isocitrate, isocitrate dehydrogenases (*Idh3b* and *Idh3g*) that form 2-oxoglutaric acid, succinate coenzyme A ligase (*Sucla2, Suclg1, Suclg2)*, succinate dehydrogenases (*Sdha, Sdhb* and *Sdhc*) that forms fumarate, fumarate hydratase 1 (*Fh1*) that produce malate, and mitochondrial malate dehydrogenases (*Mdh1 and Mdh2*) that form oxaloacetate. The pathway is derived from the KEGG database. Blue – upregulated genes, red downregulated genes, grey – not affected genes.

### Oxidative phosphorylation is downregulated

ATP for cardiac function is provided primarily by the ETC and oxidative phosphorylation, a pathway intimately connected to the TCA cycle. Located on the cristae of the IMM, four enzyme complexes of the ETC (Complexes I-IV) extract energy from the electron transporters HADH + H^+^ and QH_2_/FADH_2_ produced by β-oxidation, glycolysis, amino acid catabolism, ketone body catabolism, and the TCA cycle while generating NAD+ and FAD. The energy from TCA is used to power the export of hydrogen ions into the cristae space creating a hydrogen ion gradient. The hydrogen ion gradient is then metered through ATP synthase to drive ATP production. The entire process was significantly affected, with most DEGs downregulated (**Figure 11**).

**Figure 11.**
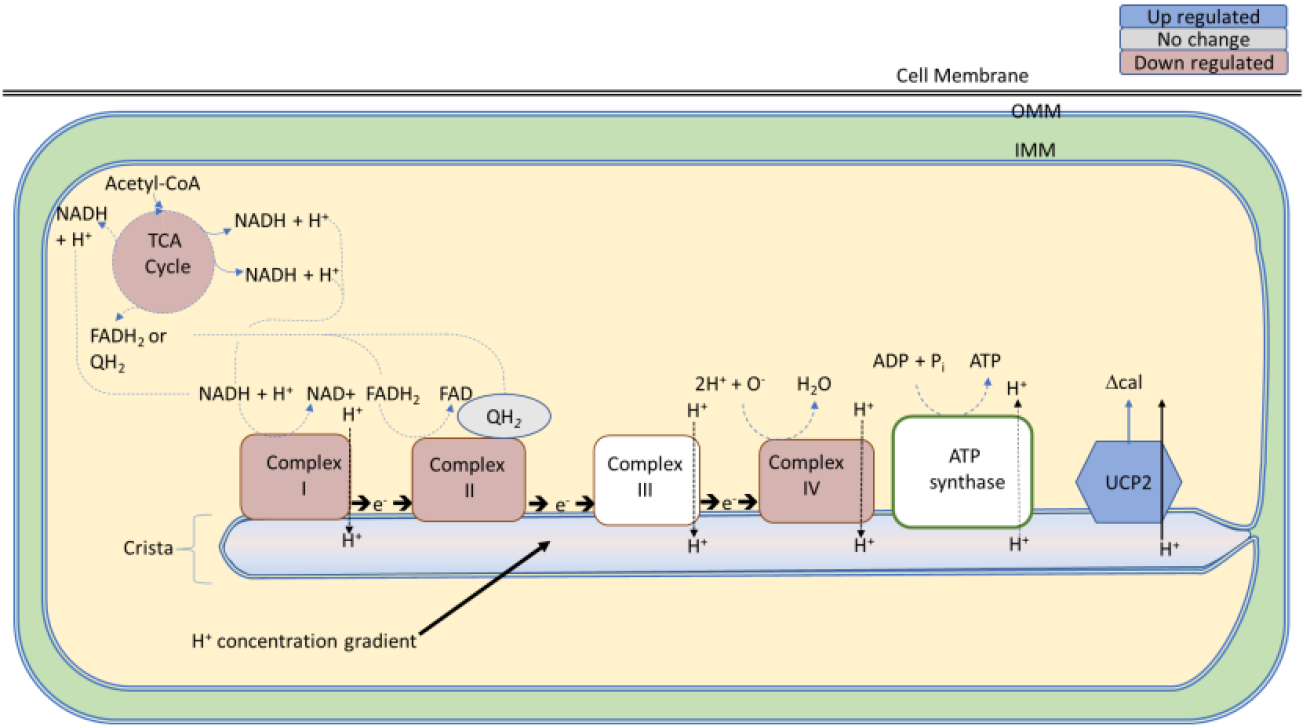
Electron transport chain and oxidative phosphorylation in oxaliplatin treated mice. Oxaliplatin affects the oxidative phosphorylation by downregulation genes associated with the proteins for Complex I, Complex II and Complex IV, reducing the strength of the hydrogen ion gradient. Uncoupling Protein 2 (UCP2 encoded by *Slc25a8*) was the only upregulated. UCP2 generate heat by dissipating the hydrogen ion gradient of the cristae instead of allowing ATP formation. The pathway is derived from the KEGG database. Blue – upregulated complexes, red downregulated complexes, grey – not affected.

Three of the seven Complex I mitochondrial genes *mt-Nd4, mt-Nd6*, and *mt-Nd4l* were downregulated along with the genes for two NADH oxidoreductase subunits (NDUFs): *Ndufaf8* and *Ndufs2* (58) (**Figure S13**). The downregulation of these genes suggests mitochondrial complex I deficiency. This would cause a wide variety of symptoms in the affected organs that require a significant amount of energy including the heart (59). In Complex II, an assembly subunit encoded by *Sdhaf4* out of eight known subunits (60) exhibited down-regulation (**Figure S14**). Complex IV had two out of 19 subunit genes: *mt-co2* and *Cox7a1*, downregulated. Interestingly, none of the genes in Complex III were affected and none of the known 25 genes that encode proteins in ATP synthase(61), exhibited statistically significant differential expressions. Overall, significant downregulation of genes that encode major complexes in the ETC suggests a decrease in hydrogen ion transport into the cristae. This suggests a decrease in ATP production in the heart. Highly correlated gene expression within ETC as shown in **Figure S14** provided a synchronized response to oxaliplatin.

One of the hallmarks of the oxaliplatin treatment in mice is the lowering of the core body temperature of animals. After 8 injections of oxaliplatin the body temperature of the 10 mg/kg oxaliplatin treated mice decreased to 33.35 ± 1.73°C (n=8, μ ± STD) compared to the control group (38.04°C ± 0.17°C, n=8) (*p*<2.54e-6, ANOVA). In response to the body temperature drop, the gene encoding the mitochondrial uncoupling protein UCP2, also known as Slc25a8, was significantly upregulated (FC=2.46). While UCP2 is known to play a role in thermogenesis and energy expenditure, it may be plausible that the increased *Ucp2* expression in the heart compensates for low core temperature. Some studies have also suggested that UCP2 protect the heart from ischemic injury by regulating mitochondrial respiration and reducing reactive oxygen species (ROS) production (70). Increased expression of this gene in response to low core temperature may protect the heart from damage by reducing oxidative stress and maintaining mitochondrial function.

### Glycolysis is upregulated

The reduced FA β-oxidation, amino acid catabolism, and ketone body catabolism in oxaliplatin-treated mice expectedly leads to decrease of ATP synthesis. The ATP deficit triggered upregulation of the *Ppargc1a* gene (FC=1.37, **Figure S15**) that encodes a master regulator energy-sensing PGC1α protein. This protein enables the heart to respond to many energy metabolism-related stimuli including cold, fasting, exercise, and changes in substrate availability (62). Upregulation of this gene in the liver is a well-established factor in gluconeogenesis activation, allowing glucose to be more readily available (63). The role of PGC1α in the heart is less known. However, the cardiac-specific overexpression of PGC1α in mouse models has been often associated with heart conditions such as dilated cardiomyopathy (64) and accelerated cardiac aging (65). The upregulation of *Ppargc1a* and its tight correlation with other upregulated genes in the glycolysis pathway, such as *Eno1* (*P*=0.94), *Gpi1* (*P*=0.93), *Pfkp* (*P*=0.93) (**Figure S16**) may suggest the central role of *Ppargc1a* for switching to glycolysis as an alternative metabolic pathway to FAs.

This switch from aerobic metabolism to anaerobic metabolism was corroborated by the upregulation of *Slc2a1* (**Figure S17)** that encodes a glucose transporter GLUT1 required to facilitate glucose transport into cardiomyocytes and the overexpression of many genes in the glycolysis pathway as illustrated in **Figure 12** and tabulated in **Table S1**. During embryonic development, GLUT1 is typically expressed in the heart and plays a vital role in glucose transport. However, at the postnatal stage, GLUT4 takes over as the primary regulator of glucose transport, and the significance of GLUT1 diminishes. In response to chronic insults, GLUT1 re-expresses in the heart under hypertrophic stress representing an adaptive response (66). The entire glycolytic pathway that needs to produce ATP was positively affected by oxaliplatin with most of genes upregulated (i.e., *Hk1*, FC=1.43, *Gpi1*, FC=1.27, *Pfkp*, FC=2.2, *Pgk1*, FC=1.27, *Eno1*, FC=1.74, *Pgam1*, FC=1.45, *Ldha, F*C=2.08) (**Figure S17**). Not surprisingly, the genes that encode enzymes that catalyze the reverse reactions, i.e., *Fbp2*, encoding enzyme FBP2 that hydrolyzes fructose-1,6-BP to fructose-6P, were downregulated FC=−1.95). Altogether, the activation of the energy-sensing mechanism, upregulation of many genes in the glycolysis pathway together with the downregulation of FAs oxidation processes suggests a shift to glycolysis to compensate for ATP generation.

**Figure 12.**
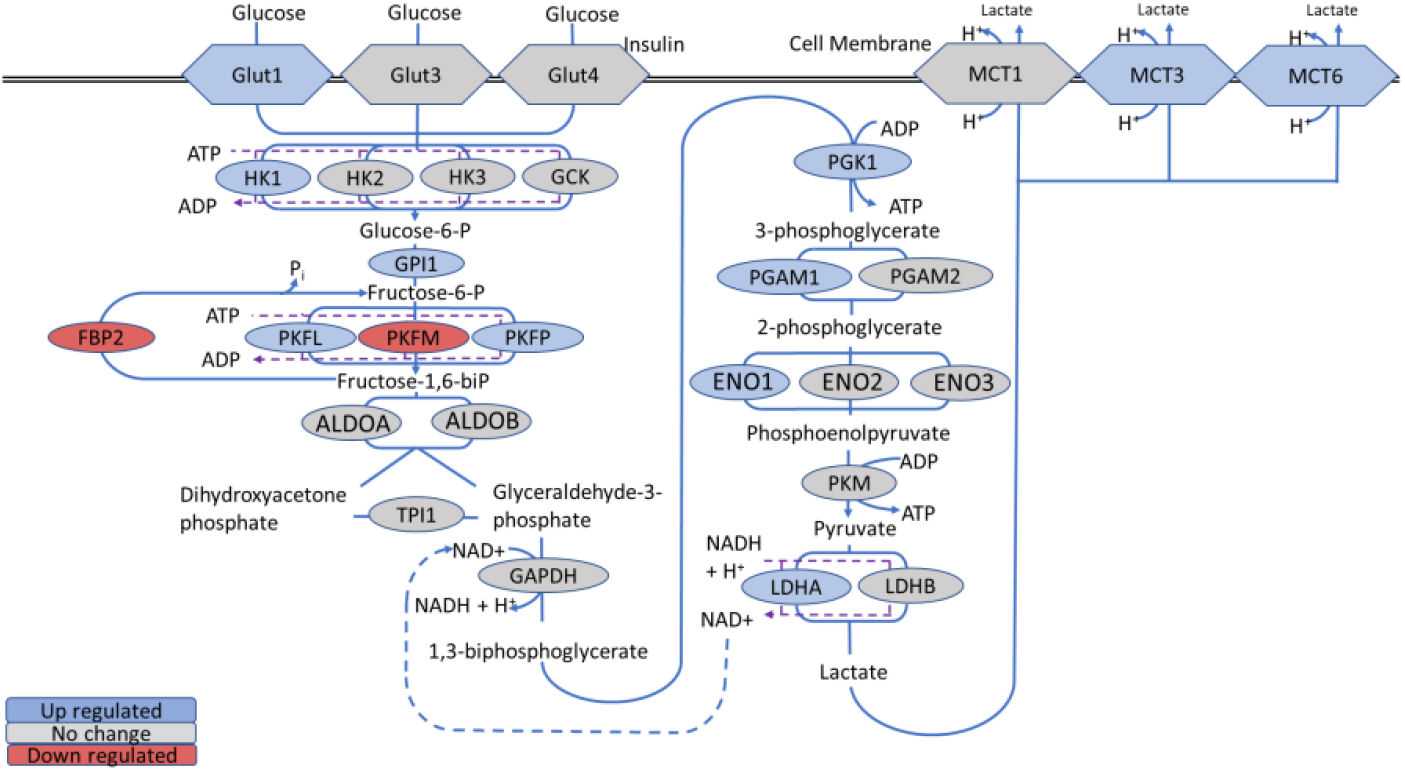
Glycolysis in cardiomyocytes is upregulated in the oxaliplatin treated mice. Abbreviations of enzymes and proteins with encoded genes is in parenthesis: ALDOA, aldolase A (*Aldoa*); ALDOB, aldolase B (*Aldob*); ENO1/2/3, enolase 1, 2 and 3 (*Eno1, Eno2* and *Eno3*); GAPDH, glyceraldehyde-3-phosphate dehydrogenase (*Gapdh*); GCK, glucokinase (*Gck*); GLUT1, glucose transporter 1 (*Slc2a1*); GLUT3, glucose transporter 3 (*Slc2a3*); GLUT4, glucose transporter 4; HK2, hexokinase 2/3 HK2 and HK3 (*Hk2* and *Hk3*); LDHB, lactose dehydrogenase B *(Ldhb*); PGAM2, phosphoglycerate mutase 2 (*Pgam2*); MCT1,3,6 monocarboxylic acid transporters 1, 3, 6 (*Slc16a1, Slc16a3, Slc16a6*); phosphofructokinases, PKFL, liver type (*Pfkl*); PKFM, muscle type (*Pfkm*); and PKFP, pallet type (*Pfkp*); PKM, pyruvate kinase, muscle type (*Pkm*); TPI1, topoisomerase 1 (*Tpi1*). The schematics is in part based on KEGG Pathways and ref (67). Blue – upregulated genes, grey – unaffected genes, red downregulated genes.

### Lactate production is upregulated

ETC malfunctioning due to oxaliplatin treatment prevents this pathway from using the power of electrons from NADH + H+ and QH2/FADH2 to transport hydrogen ions into the mitochondrial intermembrane space leading to a decrease in ATP synthesis via oxidative phosphorylation. At this point, the cell would be forced to convert the product of glycolysis, pyruvate, to lactate by using the energy from oxidizing NADH to NAD^+^ so the cell can continue the production of ATP via glycolysis (68). Indeed, the *Ldha* gene that encodes one of two subunits of lactate dehydrogenase (LDH) that converts pyruvate to lactate (69) was significantly upregulated (FC=2.08) (**Figure S18**). Similarly, two of the genes that encode the lactate transporter MCT3 and monocarboxylic acid transporter MCT6 responsible for exporting lactic acid from the cytoplasm to blood (70, 71), were also upregulated (FC=3.27 for MCT3 and FC=1.62 for MCT6). The lactate that was exported from the cardiomyocytes to the blood can then be processed by the liver via the Cori cycle to produce the needed ATP. Such upregulation in the lactate producing pathway was in sync with glycolysis activation and downregulation of ETC and may lead to lactate accumulation.

### Nicotinamide dinucleotide synthesis is activated

Among 275 DEGs from KEGG-defined metabolic pathways, *Nmrk2* from the NAD+ pathway (**Figure 13**) was the top upregulated gene (FC = 16.44) (**Figure S19**). *Nmrk2* in mice is primarily expressed in cardiomyocytes and myocytes in the skeleton muscle (72). Located in the cytosol, *Nmrk2* encodes nicotinamide riboside kinase 2 (NRK2) that phosphorylates nicotinamide riboside (NR), a derivative of vitamin B3, to generate nicotinamide mononucleotide (NMN), an immediate precursor of NAD^+^. Interest in *Nmrk2* has recently increased due to its strong connection to cardiac-related diseases and aging (73). Similar high increases in *Nmrk2* expression (FC=16.87) have been reported in glycogen synthase kinase 3 deficient hearts (74) and in mice models of dilated cardiomyopathy (73, 75). Interestingly, a shortened splice variant form of NRK2 termed the muscle integrin binding protein (MIBP) binds to the integrin β1 subunit of the heterodimer integrin α7β1 but lacks nicotinamide riboside kinase activity (73). Binding of β1 leads to disruption of laminin deposition to the lamina lucida matrix of the basement membrane of cardiac muscle cells (73, 76). This disruption of the basement membrane of cardiomyocytes may be a significant contributing factor to the remodeling observed during oxaliplatin treatment.

**Figure 13.**
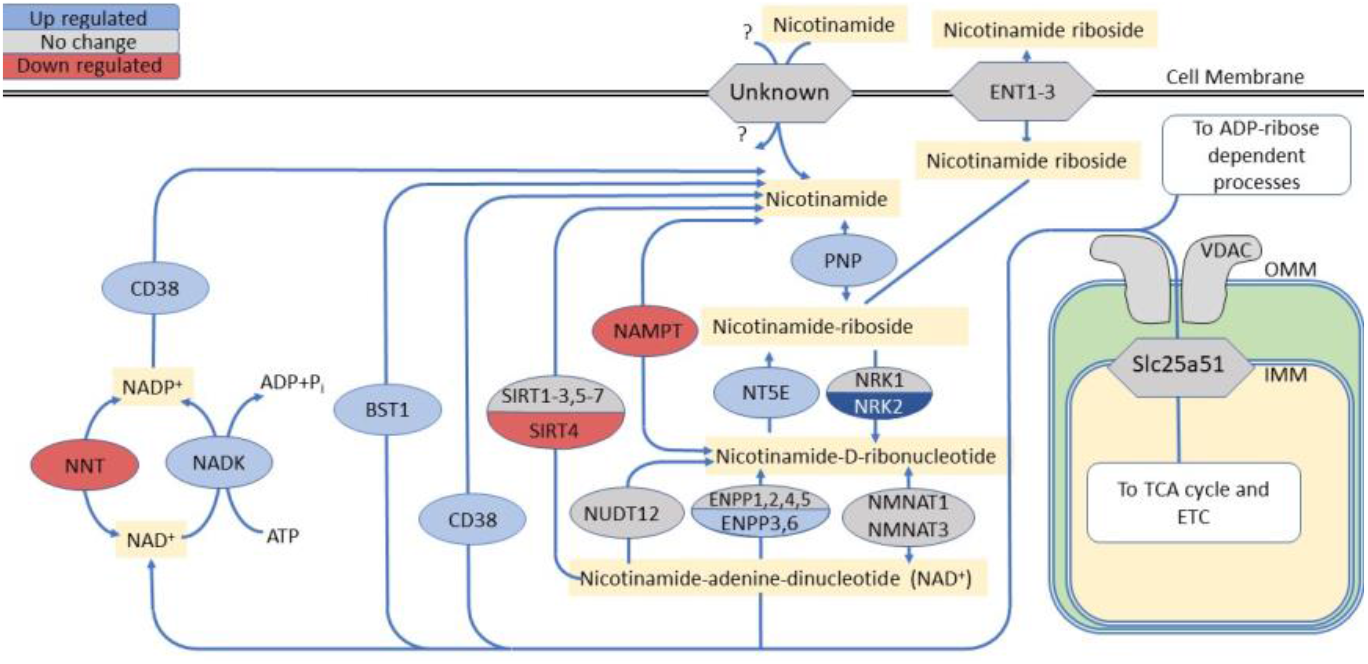
Effect of oxaliplatin on NAD+ and NADP+ synthesis and salvage pathways. The overall upregulation of the genes point to increased attempts to rebuild the cellular NAD+ pool. **Abbreviations:** CD38, NAD+ nucleosidase (*Cd38*); ENPP 1/2/3/4/5/6, ectonucleotide pyrophosphatase /phosphodiesterase 1-6 (*Enpp1, Enpp2, Enpp3, Enpp4, Enpp5, Enpp6*); NADK, NAD kinase (*Nadk*); NAMPT, nicotinamide phosphoribosyltransferase (*Nampt*); NNT, nicotinamide nucleotide transhydrogenase (*Nnt*); NRK2, nicotinamide riboside kinase 2 (*Nmrk2*); NT5E, ecto 5’-nucleotidase or CD73 (*Nt5e*); PNP, purine nucleoside phosphorylase (*Pnp*); ENT1, equilibrative nucleoside transporters 1/2/3 (*Slc29a1, Slc29a2, Slc29a3*); MCT1, monocarboxylic acid transporter 1 (*Slc16a1*); NADSYN1 (NAD Synthetase 1, *Nadsyn1*); NMNAT1/3 (nicotinamide nucleotide adenylyltransferases 1 and 3 (*Nmnat1* and *Nmnat3*); NRK1 (nicotinamide riboside kinase 1 (*Nmrk1*); NUDT12; nudix hydrolase 12 (*Nudt12*); SIRT1/2/5/6 (sirtuins, *Sirt1, Sirt2, Sirt5*, and *Sirt6*); Slc25a51, mitochondrial nicotinamide adenine dinucleotide transporter (*Slc25a51*). ‘?’ - transporter for nicotinamide remains unidentified in heart. The pathway is derived from the KEGG database. Blue – upregulated genes, red downregulated genes, grey – not affected genes.

Other genes in the NAD^+^ pathway were also upregulated (**Figure 13**), however, none of them had as large a fold change as *Nmrk2*. The extensive upregulation of *Nmrk2* in oxaliplatin-treated mice, the heart specificity, and the overall high level of copy number make this gene and its encoded enzyme attractive candidates for the assessment of oxaliplatin-induced heart damage.

*Nmrk2* expression was positively correlated with *CD38*, (P=0.71) (**Table 2**). That is not surprising given that NRK2 and CD38 are part of the same metabolic pathway for NAD+ synthesis. Their expression levels are tightly regulated to maintain NAD+ homeostasis. Higher expression of CD38 leads to decreased NAD+ levels while a higher expression of NRK2 may be required to regenerate NAD+. The tight coordination of *Nmrk2* and *CD38* expression is necessary to maintain proper NAD+ levels and cellular function. Interestingly, *Nmrk*2 expression also strongly correlated with many genes across the glycolytic pathway suggesting a synchronized response to oxaliplatin. For example, *Ppargc1a* (energy sensor and the master regulator of mitochondrial biogenesis), *Eno1* (potential hypoxia sensor (77)), *Gpi1* and *Hk1* (enzymes for glycose conversion), *Ldha* (the enzyme that converts pyruvate and NADH to lactate and NAD+), *Pfkp* and *Pfkl* (key players in glycolysis regulation) all showed strong correlations with *Nmrk2* with the corresponding *P* values above 0.75. Low negative correlation between *Nampt* and *Nmrk2* in oxaliplatin-treated mice was also expected, given that two genes often show negative correlation in murine models of cardiomyopathy and human heart failure (75).

**Table 2.**
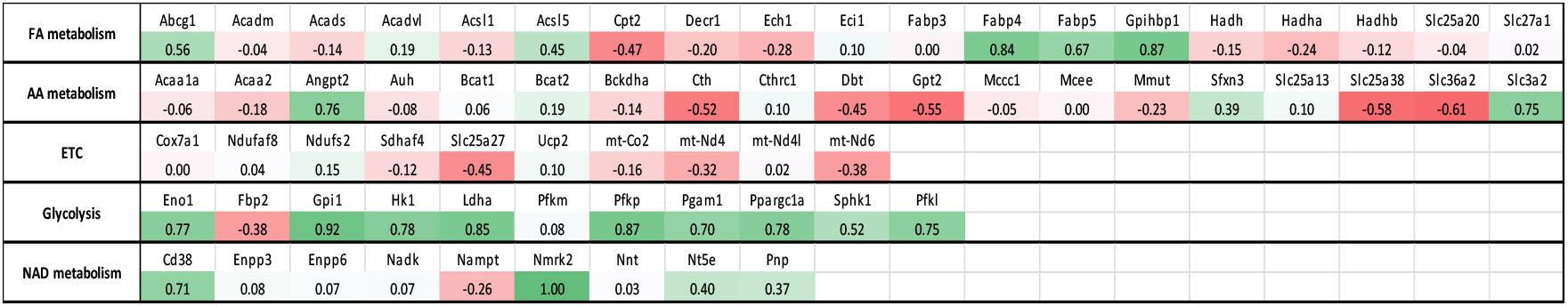
Pearson correlation coefficients of *Nmrk2* with differentially expressed genes in the energy metabolic pathway.

#### Comparison with database on human heart disease

To determine whether the changes in cardiac gene expression after oxaliplatin treatment correspond to those in common cardiac disorders, DEGs were overlaid with published human data from patients with end-stage dilated cardiomyopathy (DCM) (78). This dataset was generated from a microarray analysis and identified 129 genes co-regulated with our dataset. Among these genes are known markers associated with heart pathology such as fibronectin 1 (*Fn1*) (79), periostin (*Postn*) (80), natriuretic peptide B (*Nppb*) (81), four and a half LIM domains 1 (*Fhl1*) (82), and cyclin-dependent kinase 1 (*Cdk1*) (83). Even higher levels of genetic overlap were found between the DEGs from oxaliplatin treated mice and the DEGs from RNA-seq of a dataset of human heart failure composed of dilated cardiomyopathy (DCM) and ischemic cardiomyopathy (ICM) human heart samples (84). Total, 460 genes were similarly differentially expressed (FDR < 0.05) (**Figure 14**) with many genes shared between our data (**Figure 14, Table 3** inside). Among the affected genes related to the energy metabolism, a number of genes in the NAD+ synthesis pathways were upregulated including *Nmrk2* (FC =1.43 for DCM and FC = 1.57 for ICM). This overlap in gene expression suggests that oxaliplatin treatment leads to significant transcriptional changes that share strong similarities between the hearts from mice treated with oxaliplatin and human heart failure samples. Overall, these results support the notion that oxaliplatin may cause or aggravate preexisting heart conditions.

**Table 3.**
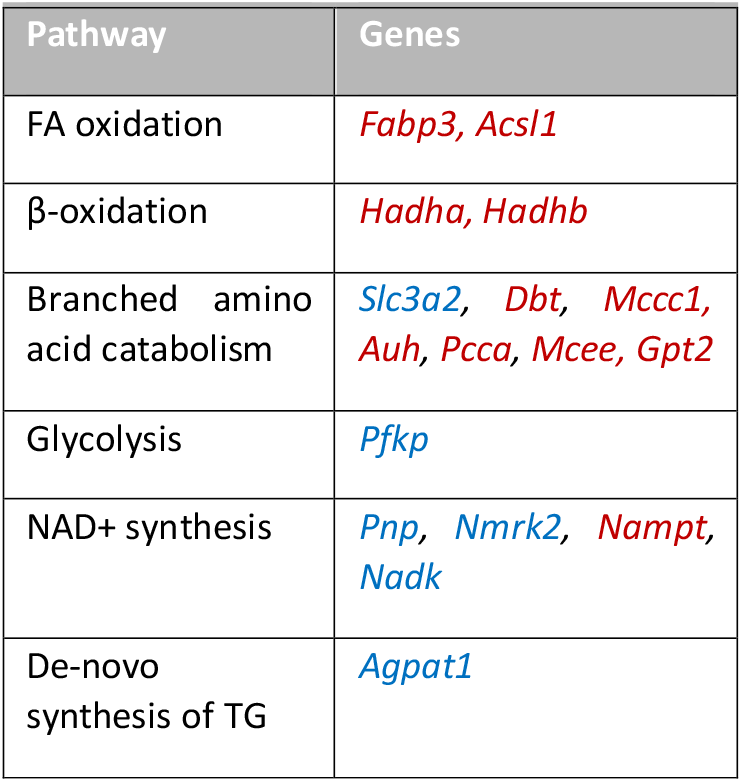
From human energy metabolism genes

**Figure 14.**
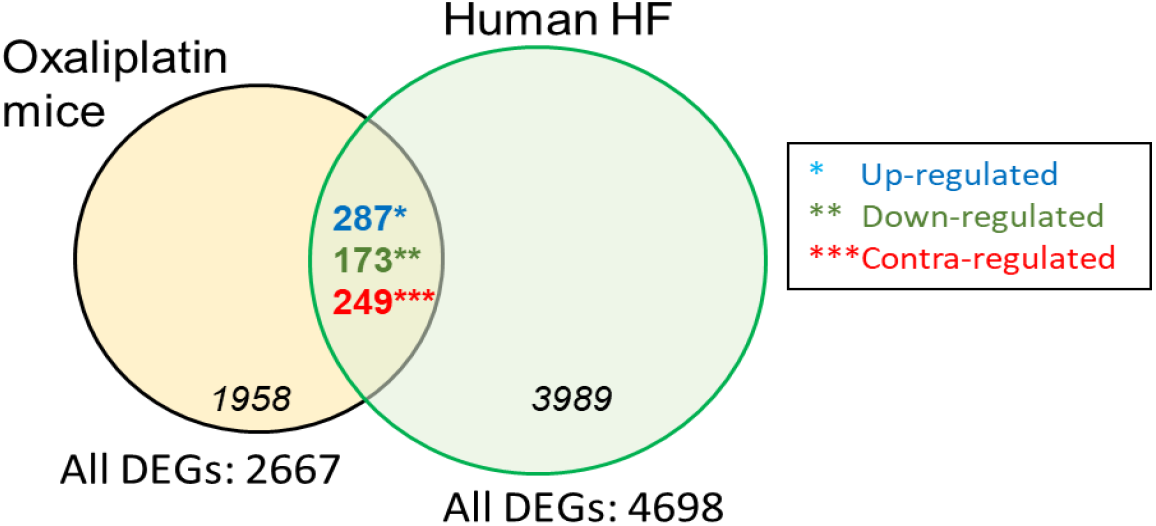
Venn diagram of DEGs in oxaliplatin treated mice vs human heart failures from dilated (DCM) and ischemic (ICM) heart samples (left ventricles). Out of 460 commonly DEGs, 16 genes were from the energy-related metabolic pathways. Blue – upregulated genes, red downregulated genes, green – contra-regulated

## DISCUSSION

The heart is highly adaptable and can easily switch from one source of energy to another when required. This metabolic flexibility allows the heart to increase its tolerance to the toxic stimuli in the short term. However, chronic metabolic insult can disrupt the normal metabolic pathways and lead to the accumulation of toxic metabolites, oxidative stress, inflammation, and cell death. These processes can result in remodeling of the blood vessels and structural changes in the heart, including hypertrophy, inflammation, and fibrosis.

The cumulative oxaliplatin dose appears to be the most significant risk factor for the development of oxaliplatin cardiotoxicity. No significant changes in the heart activity of mice by ECG receiving a human-equivalent dose of oxaliplatin was observed in the first 4 weeks of the treatment. The heart rates were significantly decreased from a normal 370 – 460 bpm range (20) to 200-370 bpm at the high-accumulated dosage, while the size of the heart became larger. After 8 weeks of treatment, the histology of the heart also revealed a number of lesions evidencing moderate to severe necrosis and inflammatory cell infiltration.

The observed cardiovascular changes from the oxaliplatin chemotherapy project a complex interaction among hemodynamic, inflammatory, electrophysiological, and neuronal circuits on the heart cells, affecting a great number of pathways. To reflect this complex interplay, we utilized a whole heart RNA-seq. The differential expression analysis presented a strong impact of oxaliplatin on the energy metabolic pathways that includes decline of FA oxidation, slowness of the TCA cycle, decrease ETC efficiency, upregulation of glycolysis with a shift to lactate production and changes in NAD+ synthesis. The multiple effects of oxaliplatin on the energy metabolic pathways are schematically summarized in **Figure 15**.

**Figure 15.**
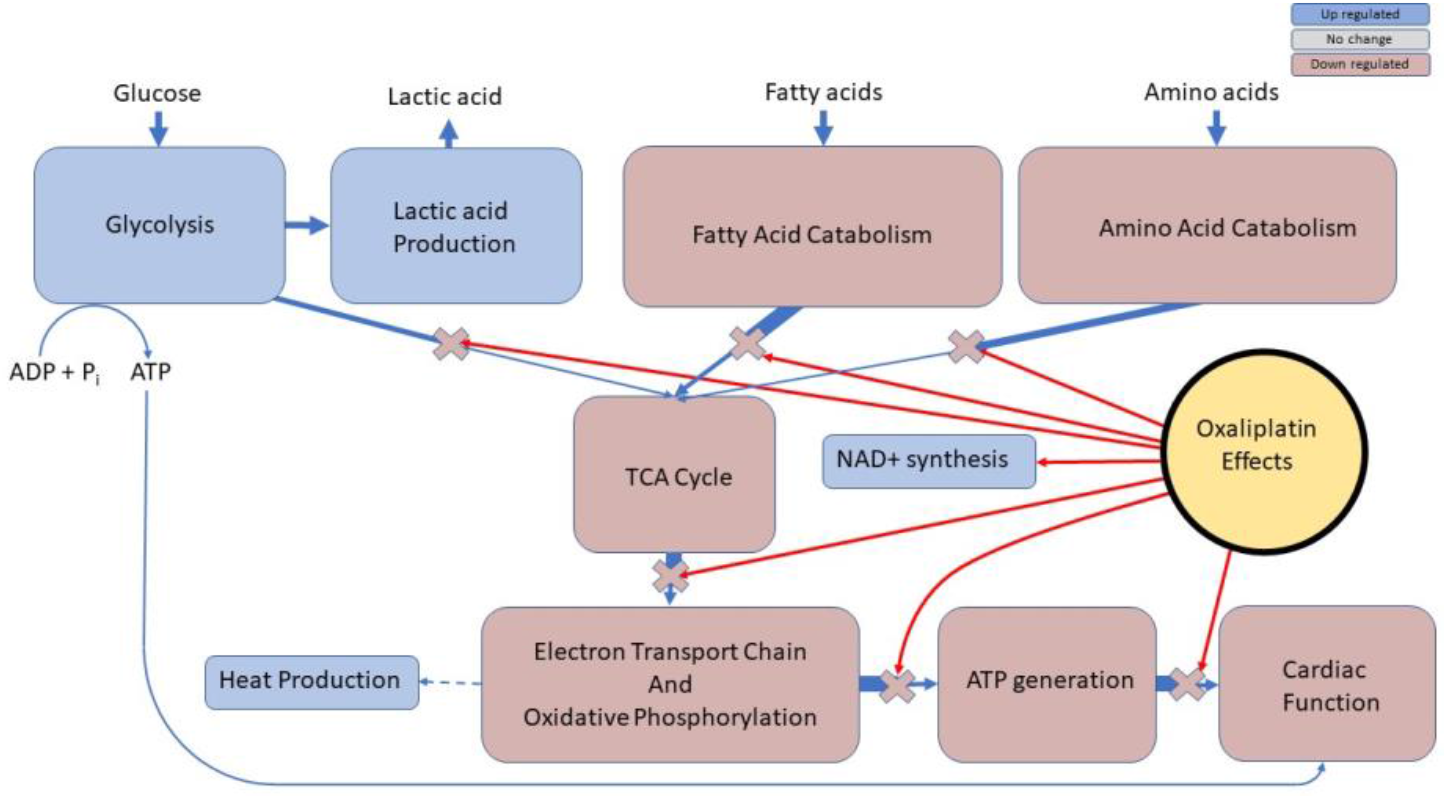
Summary schematics of oxaliplatin induced changes in the heart energy metabolic pathway. Blue – upregulated pathways, brown - downregulated pathways.

The observed energy related metabolic changes were tightly connected with other processes affected by oxaliplatin. One of the well-established effects of oxaliplatin is a strong hematological toxicity that decreases the number of red blood cells (85). Although the details of this process are not entirely understood, the low level of red blood cells may lead to the lower level of oxygen delivered to the cardiac tissue. In that regard the change from FA oxidation to glycolysis might be beneficial because anaerobic glucose metabolism is more ‘oxygen sparing’, with a complete change from FA to glucose metabolism reducing oxygen demand by 11–13% (86, 87). This process known as the “glycolysis switch” is common in mammals and refers to the phenomenon observed in the heart under stress conditions.

While this glycolysis switch initially provides the heart with increased energy production, sustained activation of glycolysis can cause metabolic dysfunction and contribute to heart failure. Specifically, chronic reliance on glucose metabolism can lead to detrimental effects on cardiac function. This is because the heart is not well adapted to rely solely on glucose metabolism and is not able to efficiently utilize glucose for energy production in the long term. Chronic glycolysis leads to the accumulation of toxic intermediates and oxidative stress, and the upregulation of glycolytic enzymes can lead to altered calcium handling and impaired contractile function in the heart. Indeed, RNA-seq showed changed expression levels of known markers of the heart damage outside of the metabolic pathway *Nppb, Myh6, Col3a1*, and *Mmp2* (**Figure S20**) (88-91).

NADH produced during glycolysis is normally re-oxidized to NAD+ through the ETC channel, but this process requires oxygen and a well-functioning ETC. In the absence of oxygen (due to the lack of blood cells) and the observed downregulated ETC, the NADH generated during β-oxidation and glucose metabolism is not being re-oxidized, leading to a decrease in the NAD+/NADH ratio. This decrease is considered detrimental on the heart as it impairs the function of enzymes that depend on NAD+ by using it as substrates or co-factors (92, 93).

Maintaining the NAD+ pool is crucial for the proper functioning of the heart. This is especially important during conditions where the heart undergoes a long-term switch to glycolysis as a means of generating ATP. The observed large increase in *Nmrk2* expression when β-oxidation and ETC are inhibited suggests that the lack of NAD+ is compensated through a complementary mechanism. This supports the idea that the NRK2 mediated NAD+ synthesis pathway represents a common adaptive mechanism in the failing heart where NAD+ levels are low (94). Linked to the lack of food consumption and weight loss of oxaliplatin-treated mice, *Nmrk2* is strongly associated with oxaliplatin tolerance, suggesting that higher levels of NAD+ might be beneficial. While the correlation between the *Nmrk2* level and weight loss in oxaliplatin-treated mice is strong, it is possible that other factors could contribute to this relationship. The treatment of mice to increase the NAD+ level such as supplementation with nicotinamide riboside might improve heart function in oxaliplatin treatment patients if this treatment does not interfere with the major purpose of oxaliplatin-based chemotherapy to eradicate cancer.

## SUMMARY AND FUTURE WORK

There is currently no method to for the early detection of or prevention of oxaliplatin-induced cardiotoxicity. These study findings help begin to elucidate the underlying metabolic pathways in order to develop screening and treatment strategies. The increased production of lactate and changes of the level of other metabolic markers can be diagnosed with established and recent clinical imaging modalities such as SPECT/PET (95), or hyperpolarized MRI (96). New markers based on the NAD+ pathway need to be established, since conventional biomarkers such as cardiac troponins might not directly reflect the changes in the heart associated with oxaliplatin-induced cardiac toxicity. Novel cardioprotective strategies (i.e., based on NAD+ stimulators) may need to be developed. Potential targets for metabolic therapy include drugs that rebalance the pyruvate-lactate axis to augment mitochondrial oxidation. Given the increasing burden of both heart failure and cancer in the aging population, the development of new biomarkers, imaging methods and cardioprotective strategies will be essential to minimize the impact of cancer therapy– associated cardiac toxicity.

## MATERIALS AND METHODS

### Animal models

All animal studies were conducted in compliance with the Washington University Institutional Animal Studies Committee (Animal Welfare Assurance #A-3381-01) and NIH guidelines. C57BL/6 male mice 8-10 weeks old were purchased from Charles River and housed in a central animal care facility with food and water *ad libitum*. Prior to the injection, clinical grade oxaliplatin (Sandoz) was diluted in 5% dextrose. Mice were weighed and an appropriate oxaliplatin dose was delivered for that animal’s mass. The treatments consisted of control (5% dextrose), and oxaliplatin dose (10 mg/kg, accumulated dose 0-80 mg/kg). The rationale of selecting the dosage is given in the Supplemental Information. To avoid circadian effects the mice were treated at the same time of day at 2 - 3 pm. At the end of the experiments, mice were anesthetized via isoflurane induction (2% at 1.5 L O_2_/min) and euthanized by cervical dislocation.

### Electrocardiogram (ECG)

ECG analysis on each mouse was conducted biweekly to examine the heartbeats and changes in the ECG waveforms that would alert for the risk of morbidity or mortality of the mouse. ECGs were measured one week prior to the start of the experiment and 48 h after drug deliveries. Prior to ECG, mice were anesthetized via isoflurane induction (2% at 1.5 L O_2_/min). Each mouse was transferred to a battery-powered heating pad (Kent Scientific) in a grounded Faraday cage and maintained on isoflurane for the duration of the recording. Stainless steel electrodes (Fine Science Tools) were used transdermally to record the electrocardiogram. Electrodes, skin, and fur over the electrode insertion site were sanitized with isopropanol. A three-lead electrocardiogram configuration, equivalent to human V1 chest lead configuration was used. The positive electrode was inserted through the skin of the ventral chest 4 mm rostrad to the xiphoid process along the sternum, the negative electrode was inserted through the skin of the upper right chest over the pectoralis muscle. The ground electrode was inserted through the skin on the medial surface of the right thigh. An extracellular amplifier (DAM80, World Precision Instruments) was used to record the mouse ECG (gain 100x, hi pass filter 0.1 Hz, low pass filter 1 Hz). Data were collected via Powerlab 8/35 (AD Instruments) and analyzed via software (LabChart, AD Instruments).

### Histological analysis

At the completion of week 8, the hearts of the euthanized mice were dissected, and the masses were recorded. Hearts were fixed in 10% neutral buffered formalin. Tissues were trimmed and processed according to standard protocols, embedded in paraffin, and sectioned at 5 μm. Sections were stained via hematoxylin-eosin and further imaged on an Olympus microscope (BX51).

### RT-qPCR

Total RNA (200 ng) was reverse transcribed using SuperScript IV VILO Master Mix (ThermoFisher Scientific) according to manufacturer’s instructions. RT-qPCR was performed with SYBRGreen Master Mix and QuantStudio 3 Real-Time PCR system (Applied Biosystem). Ribosomal 18S was used as an internal control and relative quantification was calculated by 2−ΔΔCt method. Primer sequences for 18S, Nppb, Cdk1, and Lgals3 are as follow: 18S forward AAGTTCCAGCACATTTTGCGAGTA, reverse TTGGTGAGGTCGATGTCTGCTTTC; Nppb forward CTTCCTACAACAACTTCAGTGC, reverse AGGTGACACATATCTCAAGCTG; Cdk1 forward AAGTGTGGCCAGAAGTCGAG, reverse TGAGAGCAAATCCAAGCCGT; Lgals3 forward AGTTATTGTCCTGCTTCGTGT, reverse GTGAAACCCAACGCAAACAG; Myh6 forward GTTAAGGCCAAGGTCGTGTC, reverse GCCATGTCCTCGATCTTGTC.

### RNA sequencing of heart and the muscle

Hearts from euthanized animals were quickly collected and immersed in ice-cold RNA*later* solution (ThermoFisher Scientific). RNA was extracted with the magnetic bead based MaxwellRSC 48 instrument using the Maxwell RSC simply RNA Tissue kit, and RNA quality was assessed using Agilent TapeStation 4200 System. From the group of 10 mice used for RNA sequencing only samples used for RNA sequencing with an integrity number (RIN) greater than 8 were used. The sample with the low RIN was discarded. cDNA library was prepared according to manufacturer’s protocol with Clontech SMARTer method, indexed, pooled, and sequenced on an Illumina NovaSeq 6000. RNA-seq reads are aligned to the Ensembl release 76 primary assembly with STAR version 2.5.1a. Gene counts were derived from the number of uniquely aligned unambiguous reads in the Partek Flow software package (Partek Inc.). Data were then normalized using median ratio and DEGs were identified with DESeq2 package implemented in Partek Flow.

Hierarchical clustering analysis was performed using the following parameters: Cluster distance metric: Average Linkage; Point distance metric: Euclidean; Cluster distance metric: Average Linkage.

The results of RNA-seq will be uploaded in the Gene Expression Omnibus (GEO) and available upon acceptance of the mansucript

### Correlation analysis

Correlation analysis was performed using Matlab custom GUI (to be uploaded on Github) and using open-source data visualization package for the statistical programming language R, R studio for Windows. The correlation matrix was visualized with ggplot2 package.

### Network analysis

All statistically enriched terms using GO and KEGG pathway were performed on all 2,667 DEGs. Terms with a p-value < 0.01, a minimum count of 3, and an enrichment factor > 1.5 (the enrichment factor is the ratio between the observed counts and the counts expected by chance) are collected and grouped into clusters based on their membership similarities. More specifically, p-values were calculated based on the cumulative hypergeometric distribution (97), and q-values are calculated using the Benjamini-Hochberg procedure to account for multiple testings (98). Kappa scores (99) were used as the similarity metric when performing hierarchical clustering on the enriched terms, and sub-trees with a similarity of > 0.3 are considered a cluster. The most statistically significant term within a cluster is chosen to represent the cluster.

To further capture the relationships between the genes, a subset of enriched genes were selected and rendered as a network plot, where terms with a similarity > 0.3 are connected by edges. The network was visualized with Cytoscape (100) with “force-directed” layout and with edge bundled for clarity. Only physical interactions in STRING (physical score > 0.132) and BioGrid were used to extract protein-protein interactions to form network. The resultant network contains the subset of proteins that form physical interactions with at least one other member in the list. The Molecular Complex Detection (MCODE) algorithm (101) was then applied to this network to identify neighborhoods where proteins are densely connected (clusters). Finally, GO enrichment analysis was applied to each MCODE network to extract “biological meanings” from the network component, where top three best p-value terms were retained.

## ACKNOWLEDGEMENT

The authors thank the NCI/ NIH (R01CA208623 MB and R21CA269099 MB) for funding this study. Blood analysis has been performed by DCM at Washington University. Heart sections were examined by Dr. Suellen Greco. We thank the Alvin J. Siteman Cancer Center at Washington University School of Medicine and Barnes-Jewish Hospital in St. Louis, MO. and the Institute of Clinical and Translational Sciences (ICTS) at Washington University in St. Louis, for the use of the Genome Technology Access Center, which provided RNA-seq service. The Siteman Cancer Center is supported in part by an NCI Cancer Center Support Grant #P30 CA091842 and the ICTS is funded by the National Institutes of Health’s NCATS Clinical and Translational Science Award (CTSA) program grant #UL1 TR002345.

## Abbreviations

CIPN: chemotherapy induced peripheral neuropathy
ECG: electrocardiogram
ETC: electron transport chain
OMM: outer mitochondrial membrane
IMM: inner mitochondrial membran
(DEGs): differentially expressed genes
DCM: dilated cardiomyopathy
ICM: ischemic cardiomyopathy
FAs: fatty acids
TCA: tricarboxylic acid cycle

