## Supplemental File for "Oxaliplatin-induced cardiotoxicity in mice is connected to the changes in energy metabolism in the heart tissue"

### Table S1 Summary of the metabolic genes affected by oxaliplatin.

| Name | Abbreviation | Gene | Fold change (FC) | FDR | LS mean oxaliplatin | LS mean control |
|---|---|---|---|---|---|---|
| **1. FA OXIDATION PATHWAY** | | | | | | |
| ***1a. FAs transport from capillary lumen to endothelium*** | | | | | | |
| Glycosylphosphatidylinositol-anchored protein 1 | GPIHBP1 | *Gpihbp1* | 2.51 | 8.56E-5 | 13,048 | 5,201 |
| FA binding protein 3, Heart FA binding protein | H-FABP | *Fabp3* | -1.44 | 9.45E-3 | 151,309 | 217,409 |
| FA binding protein 4 | FABP4 | *Fabp4* | 2.01 | 2.41E-4 | 34,649 | 17,215 |
| FA binding protein 5 | FABP5 | *Fabp5* | 3.12 | 2.79E-10 | 3,366 | 1,080 |
| Solute carrier family 27 (FA transporter), member 1 | FATP1 | *Slc27a1* | -1.53 | 8.03E-3 | 3,617 | 5,530 |
| Acyl-CoA synthetase long chain family member 1 | ACSL1 | *Acsl1* | -1.3 | 0.01 | 11,898 | 15,513 |
| Acyl-CoA synthetase long chain family member 5 | ACSL5 | *Acsl5* | 1.55 | 4.7E-6 | 485 | 312 |
| ATP binding cassette transporter g1 | ABCG1 | *Abcg1* | 2.29 | 3.01E-6 | 1,182 | 517 |
| **1b. *FA inner mitochondrial membrane transport*** | | | | | | |
| Carnitine acylcarnitine translocase | CACT | *Slc25a20* | -1.43 | 4.99E-3 | 6,423 | 9,164 |
| Carnitine palmitoyltransferase 2 | CPT2 | *Cpt2* | -1.71 | 8.6E-12 | 6,484 | 11,062 |
| **1c. *β* oxidation** | | | | | | |
| Acyl-CoA dehydrogenase short chain | SCAD | *Acads* | -1.47 | 4.55E-4 | 3,230 | 4,738 |
| Acyl-CoA dehydrogenase medium chain | MCAD | *Acadm* | -1.43 | 2.23E-3 | 22,789 | 32,507 |
| Acyl-CoA dehydrogenase very long chain | VLCAD | *Acadvl* | -1.31 | 0.05 | 51,296 | 67,421 |
| Hydroxyacyl- CoA dehydrogenase | HADH | *Hadh* | -1.5 | 2.93E-4 | 15,623 | 23,387 |
| Hydroxyacyl- CoA dehydrogenase trifunctional multienzyme complex subunit alpha | HADHA | *Hadha* | -1.53 | 2.5E-6 | 21,896 | 33,443 |
| Hydroxyacyl- CoA dehydrogenase trifunctional multienzyme complex subunit beta | HADHB | *Hadhb* | -1.47 | 1.9E-3 | 55,322 | 81,571 |

| | | | | | | |
|---|---|---|---|---|---|---|
| ***1d. Unsaturated FA β oxidation*** | | | | | | |
| Enoyl-CoA delta isomerase 1 | ECI1 | *Eci1* | -1.49 | 0.02 | 8,555 | 12,772 |
| 2,4-Dienoyl-CoA reductase 1 | DECR1 | *Decr1* | -1.44 | 3.49E-4 | 5,481 | 7,904 |
| Enoyl-CoA hydratase 1 | ECH1 | *Ech1* | -1.78 | 1.89E-6 | 53,358 | 95,125 |
| **2. AMINO ACID (AA) CATABOLISM** | | | | | | |
| ***2a. Branched AA catabolism*** | | | | | | |
| 4F2 Cell-surface antigen heavy chain | 4F2HC | *Slc3a2* | 1.40 | 0.05 | 1,365 | 974 |
| Branched chain aminotransferase 1, cytoplasmic | BCAT1 | *Bcat1* | 2.93 | 1.72E-5 | 95 | 33 |
| Branched chain aminotransferase 2 mitochondrial | BCAT2 | *Bcat2* | -1.33 | 0.04 | 5,739 | 7,617 |
| Branched chain α-keto-acid dehydrogenase alpha subunit | BCKDHA | *Bckdha* | -1.76 | 4.1E-6 | 5,002 | 8,805 |
| Dihydrolipoamide Branched Chain Transacylase E2 | DBT | *Dbt* | -1.76 | 5.16E-6 | 5,002 | 8,806 |
| ***Leucine catabolism*** | | | | | | |
| Isovaleryl CoA dehydrogenase | IVD | *Ivd* | -1.46 | 2.90E-04 | 668 | 1,174 |
| 3-methylcrotonyl- CoA carboxylase | MCCC1 | *Mccc1* | -1.21 | 4.17E-02 | 10,859 | 15,815 |
| 3-methylglutaconic- CoA hydratase | AUH | *Auh* | -1.63 | 3.02E-03 | 3,661 | 4,429 |
| ***Isoleucine catabolism*** | | | | | | |
| Acetyl-CoA Acyltransferase 1 | ACAA1A | *Acaa1a* | -1.35 | 0.01 | 1,323 | 2,155 |
| Acetyl-CoA Acyltransferase 2 | ACAA2 | *Acaa2* | -1.51 | 3.693E-4 | 1,956 | 2,635 |
| ***Valine catabolism*** | | | | | | |
| Propionyl- CoA carboxylase A | PCCA | *Pcca* | -1.40 | 1.49E-4 | 36,493 | 54,934 |
| Methylmalonyl-CoA Epimerase | MCEE | *Mcee* | -1.33 | 0.04 | 1,367 | 1,913 |
| Methylmalonyl- CoA mutase | MMUT | *Mmut* | -1.28 | 3.13E-3 | 2,030 | 2,701 |
| ***2b. Neutral AA catabolism*** | | | | | | |
| Proton-coupled amino acid transporter 2 | PAT2 | *Slc36a2* | -1.39 | 0.02 | 2,864 | 3,675 |
| Mitochondrial glycine transporter | Slc25a38 | *Slc25a38* | -1.61 | 3.57E-4 | 1,061 | 1,478 |
| Sideroflexin-3 | SFXN3 | *Sfxn3* | 2.42 | 2.69E-9 | 376 | 607 |
| Citrin | CITRIN | *Slc25a13* | -1.24 | 0.05 | 1,084 | 449 |
| Alanine aminotransferase 2 | GPT2 | *Gpt2* | -2.05 | 2.58E-5 | 2,334 | 2,903 |
| Cystathionine gamma-lyase | CTH | *Cth* | -2.63 | 1.5E-4 | 262 | 535 |
| | | | | | 28 | 74 |
| **3. KETONE BODY METABOLISM** | | | | | | |
| 3-Oxoacid-CoA-transferase 1 | | *Oxct1* | -1.63 | 1.72E-3 | 21,569 | 29,278 |
| **4. GLYCOLYSIS** | | | | | | |
| PPARG coactivator 1 alpha | PGC1α | *Ppargc1a* | 1.37 | 1.39E-3 | 3,778 | 2,754 |
| Solute carrier family 2 member 1 | GLUT1 | *Slc2a1* | 1.39 | 0.01 | 1,028 | 737 |
| Hexokinase 1 | HK1 | *Hk1* | 1.43 | 2.29E-3 | 6,470 | 4,516 |
| Glucose-6-phosphate isomerase 1 | GPI1 | *Gpi1* | 1.27 | 0.04 | 21,784 | 17,203 |
| Phosphofructokinase, platelet | PFKP | *Pfkp* | 2.20 | 1.20E-6 | 3,406 | 1,546 |
| Phosphofructokinase, muscle | PFKM | *Pfkm* | -1.22 | 0.04 | 23,511 | 28,651 |
| Enolase 1 | ENO1 | *Eno1* | 1.74 | 1.84E-6 | 18,084 | 10,423 |
| Phosphoglycerate mutase 1 | PGAM1 | *Pgam1* | 1.45 | 3.97E-3 | 2,452 | 1,692 |
| Fructose-bisphosphatase 2 | FBP2 | *Fbp2* | -1.95 | 4.18E-5 | 777 | 1,514 |
| Lactate dehydrogenase A | LDHA | *Ldha* | 2.08 | 7.89E-5 | 47,492 | 60,017 |
| **5. TCA Cycle** | | | | | | |
| 2-Oxoglutarate dehydrogenase | Ogdh | *Ogdh* | -1.26 | 2.88E-3 | 21,639 | 27,652 |

| | | | | | | |
|---|---|---|---|---|---|---|
| **6. OXIDATIVE PHOSPHORYLATION** | | | | | | |
| Mitochondrially encoded ND4 NADH dehydrogenase subunit 4 | MT-ND4 | *mt-Nd4* | -1.41 | 1.43E-3 | 333,013 | 469,414 |
| Mitochondrially encoded ND6 NADH dehydrogenase subunit 6 | MT-ND6 | *mt-Nd6* | -1.31 | 3.25E-3 | 55,138 | 72,343 |
| Mitochondrially encoded NADH 4L dehydrogenase | MT-ND4L | *mt-Nd4l* | -1.28 | 0.04 | 21,639 | 27,652 |
| NADH:ubiquinone oxidoreductase complex assembly factor 8 | NDUFAF8 | *Ndufaf8* | -1.49 | 0.01 | 1,089 | 1,617 |
| NADH:ubiquinone oxidoreductase core subunit S2 | NDUFS2 | *Ndufs2* | -1.27 | 0.05 | 28,920 | 36,697 |
| Succinate dehydrogenase complex assembly factor 4 | SDHAF4 | *Sdhaf4* | -1.38 | 0.02 | 2,143 | 2,962 |
| Cytochrome oxidase 2 | COX2 | *Mt-Co2* | -1.33 | 0.02 | 309,309 | 411,289 |
| Cytochrome C oxidase subunit 7A1 | COX7A1 | *Cox7a1* | -1.51 | 0.02 | 19,791 | 29,829 |
| Uncoupling protein 2 | UCP2 | *Ucp2** | 2.46 | 4.55E-6 | 7,383 | 2,998 |
| Mitochondrial Uncoupling Protein 4 | UCP4 | *Slc25a27* | -1.63 | 0.01 | 84 | 137 |
| **7. CORI CYCLE** | | | | | | |
| Lactate dehydrogenase A | LDHA | *Ldha* | 2.08 | 7.89E-5 | 62,082 | 29,856 |
| Moncarboxylic acid transporter 3 | MCT3 | *Slc16a3* | 3.27 | 1.63E-5 | 138 | 42 |
| Monocarboxylic acid transporter 6 | MCT6 | *Slc16a6* | 1.62 | 5.84E-3 | 313 | 193 |
| **8. NAD SYNTHESIS** | | | | | | |
| Nicotinamide riboside kinase 2 | NMRK2 | *Nmrk2* | 16.44 | 1.40E-3 | 8,183 | 498 |
| Ectonucleotide Pyrophosphatase/Phosphodiesterase6 | ENPP6 | *Enpp6* | 1.99 | 0.02 | 353 | 177 |
| Purine Nucleoside Phosphorylase | PNP | *Pnp* | 1.80 | 2.75E-6 | 698 | 388 |
| Ectonucleotide Pyrophosphatase/Phosphodiesterase 3 | ENPP3 | *Enpp3* | 1.61 | 0.01 | 178 | 111 |
| ADP ribosyl cyclase 1 | CD38 | *Cd38* | 1.61 | 0.01 | 754 | 469 |
| 5'-Nucleotidase Ecto | NT5E | *Nt5e* | 1.40 | 0.02 | 639 | 456 |
| NAD Kinase | NADK | *Nadk* | 1.22 | 0.03 | 987 | 810 |
| Nicotinamide Nucleotide Transhydrogenase | NNT | *Nnt* | -1.31 | 0.03 | 14,990 | 19,693 |
| Nicotinamide Phosphoribosyltransferase | NAMPT | *Nampt* | -1.36 | 1.23E-3 | 1,997 | 2,720 |
| **9. DE-NOVO SYNTHESIS of TG** | | | | | | |
| glycerol-3-phosphate acyltransferase 3 | GPAT3 | *Gpat3* | -1.52 | 1.42E-3 | 329 | 329 |
| 1-acyl-sn-glycerol-3-phosphate acyltransferase alpha | AGPAT1 | *Agpat1* | −1.32 | 6.65E-3 | 567 | 567 |
| 1-acyl-sn-glycerol-3-phosphate acyltransferase beta | AGPAT2 | *Agpat2* | −1.60 | 9.76E-4 | 859 | 859 |
| Diacylglycerol O-acyltransferase 2 | DGAT2 | *Dgat2* | −1.79 | 2.73E-4 | 5,938 | 5,938 |

**FIGURES**

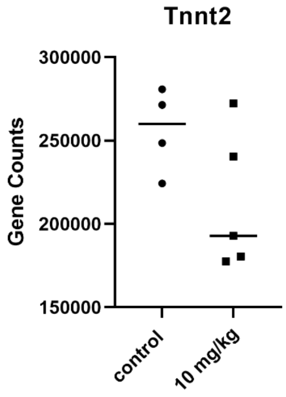

**Figure S1** Traditional blood biomarker Cardiac Troponin T (Tnnt2) is not statistically affected by oxaliplatin. FC = −1.2, FDR = 0.22.

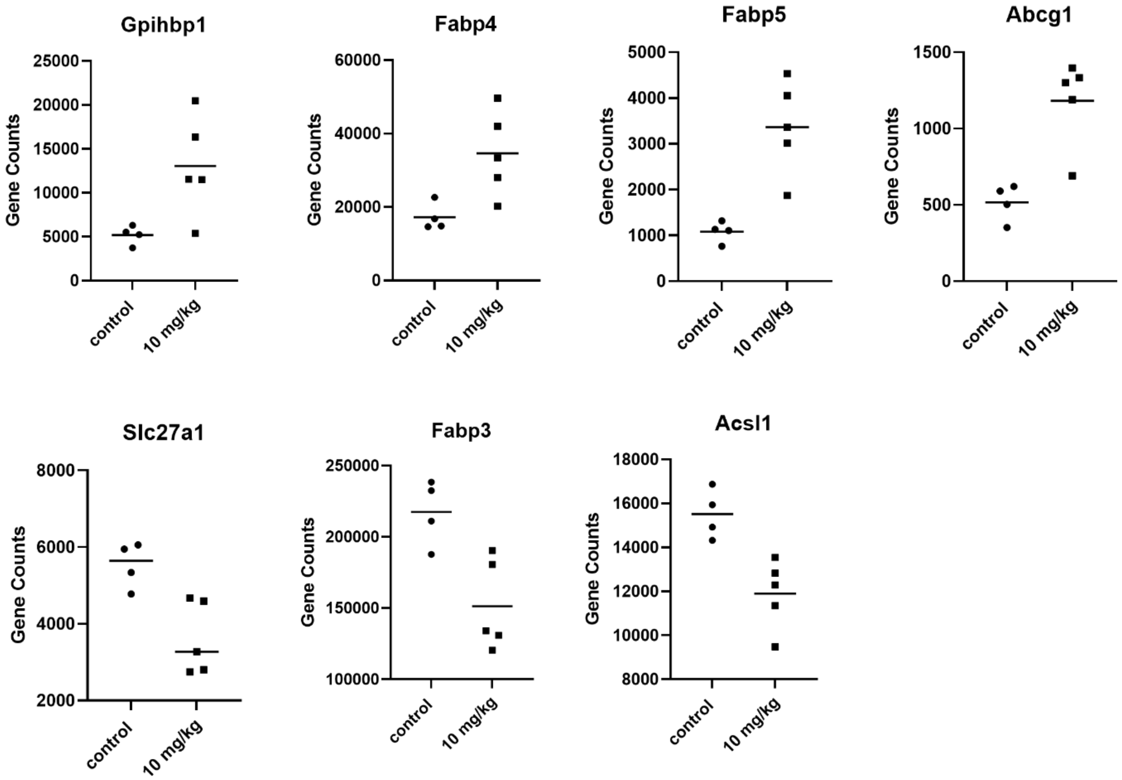

**Figure S2** Oxaliplatin upregulates genes responsible for transport of FAs from capillary lumen to endothelium: *Gpihbp1* (FC =2.51, FDR=8.56E-5), *Fabp4* (FC = 2.01, FDR=2.54E-4) and *Fabp5 (*FC= 3.12, FDR=2.79E-10), *Abcg1* (FC = 2.29, FDR = 3.01E-06). Oxaliplatin downregulates genes responsible for transport of FAs from endothelium to mitochondrion of cardiomyocytes: *Slc27a1* (FC = −1.53, FDR=8.03E-3), *Fabp3* (FC = −1.44, FDR=9.45E-3), *Acsl1* (FC = −1.3, FDR=0.02).

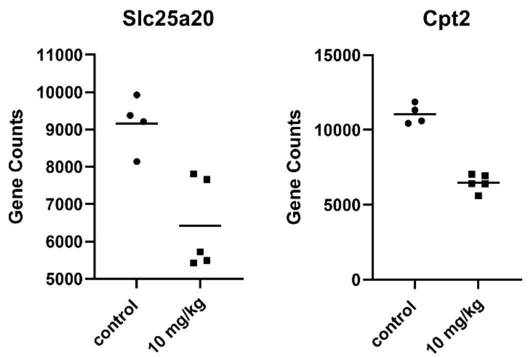

**Figure S3** Key proteins in the FA transport through mitochondria membrane are downregulated: *Slc25a20* (FC= −1.43, FDR = 4.99E-3) and *Cpt2* (FC = −1.71, FDR = 8.6E-12).

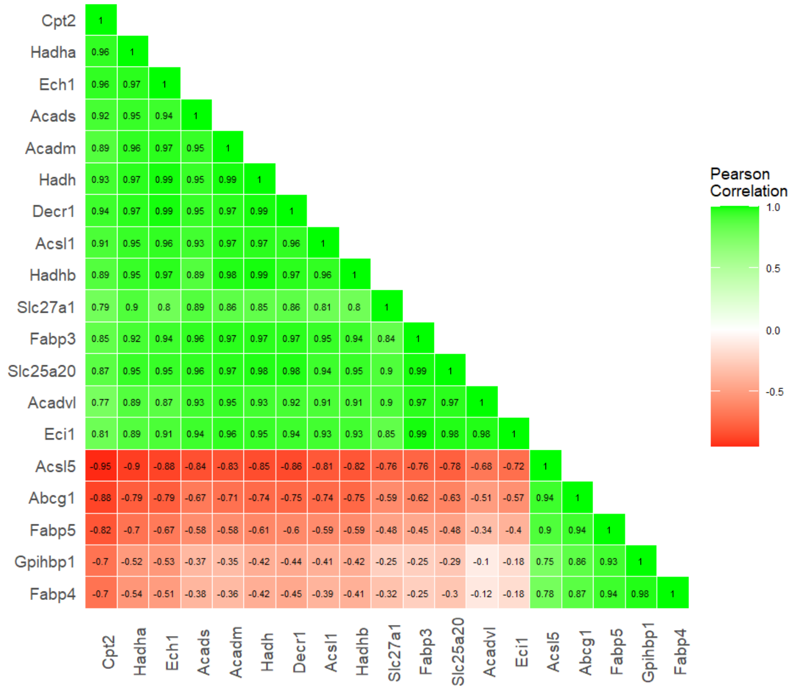

**Figure S4** FA transport and oxidation process: correlation matrix of differentially expressed genes

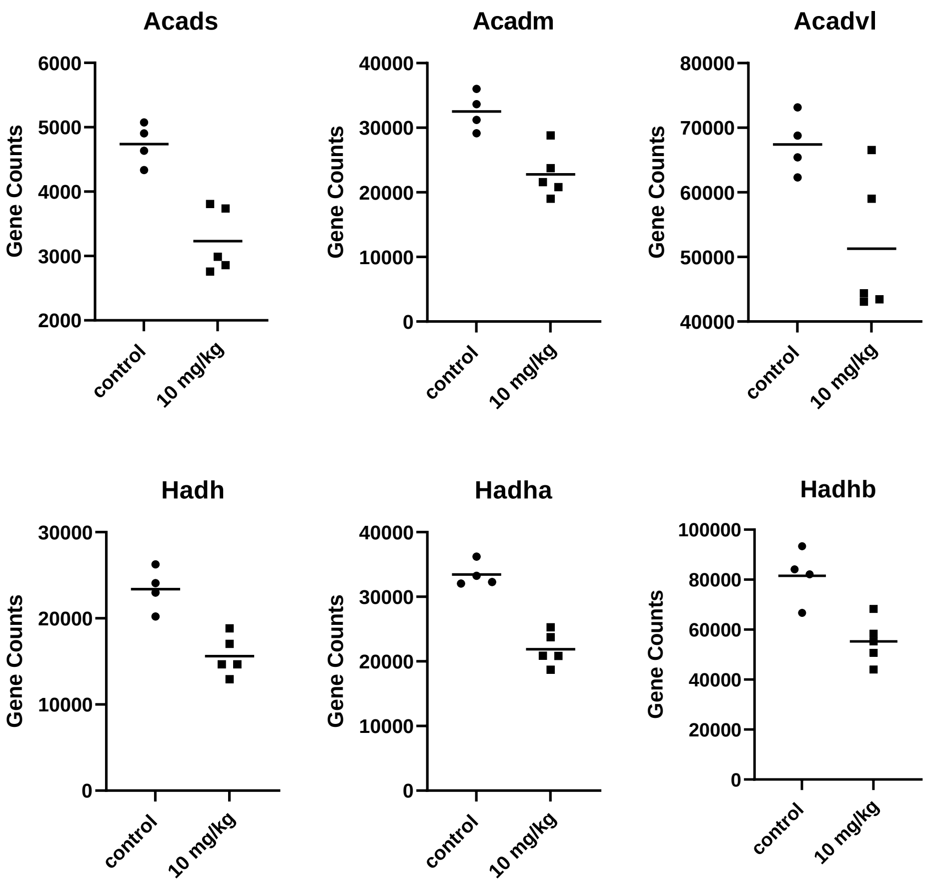

**Figure S5** Effect of oxaliplatin on β-oxidation of saturated FAs. Oxaliplatin downregulates three of the four Acyl-CoA dehydrogenases and a thiolase: short chain FA (SCAD, *Acads* (FC = −1.47, FDR=4.55E-4), medium chain FA *Acadm* (FC = −1.43, FDR=2.23E-3) and very long chain FA *Acadvl* (FC = −1.31, FDR=0.05), that corresponds to short, medium, and very long FAs. Dehydrogenation enzymes and thiolysis enzyme are also downregulated: NADH, encoded by *Nadh* (fold change= −1.5, FDR = 2.93E-4), *Nadha* (fold change= −1.53, FDR = 2.5E-6), *Nadhb* (FC = −1.47, FDR 1.9E-3).

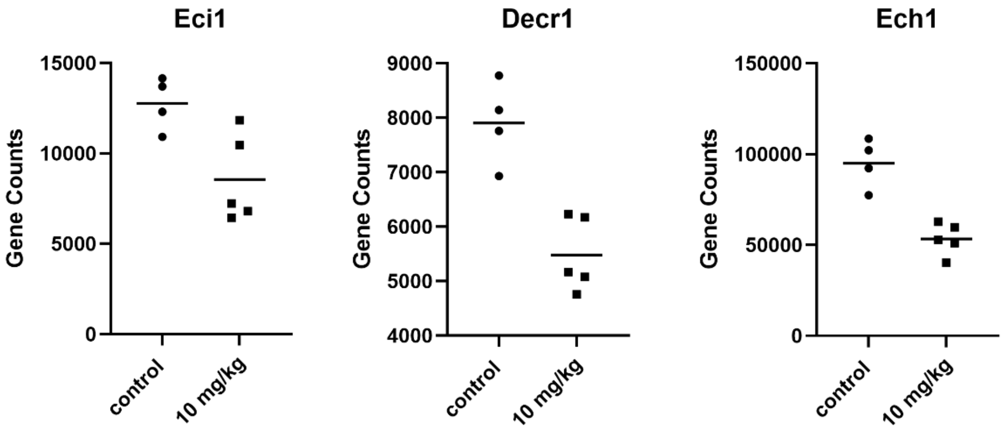

**Figure S6** Effect of oxaliplatin on β-oxidation of unsaturated FAs results on downregulation of several key genes: *Eci1* (FC = −1.49, FDR = 0.02); *Decr1* (FC = −1.44, FDR = 3.49E-4), *Ech1* (FC = −1.78, 1.89E-6)

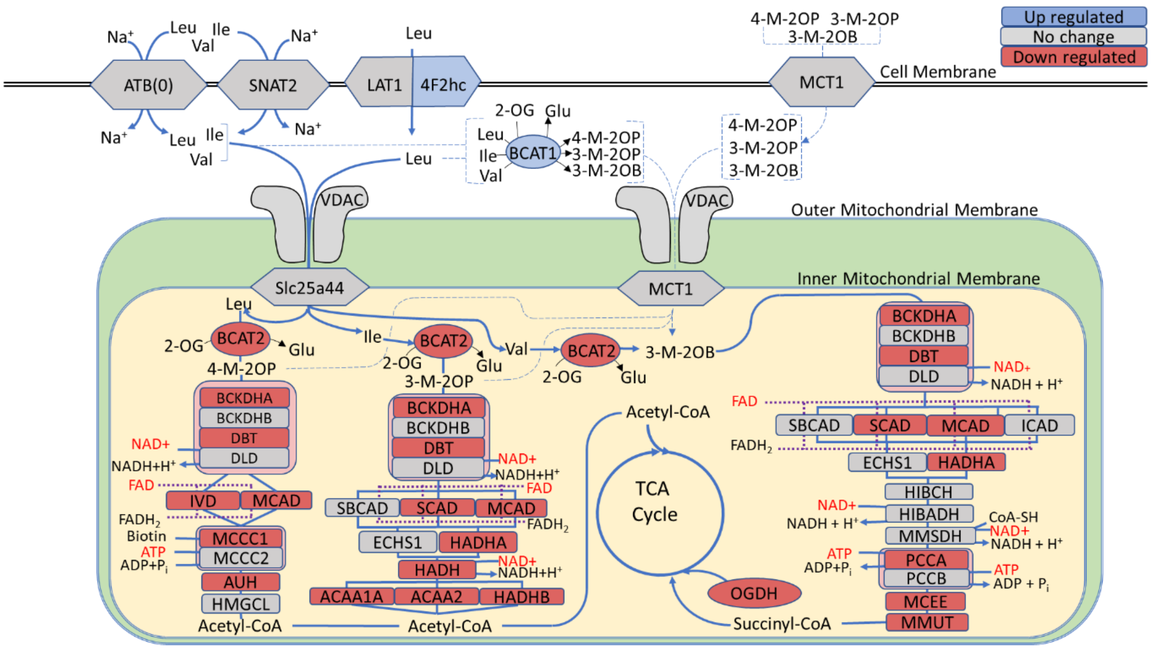

**Figure S7 Branched Amino acid metabolism is largely downregulated by oxaliplatin.** The schematics are partially based on the KEGG Pathway database.

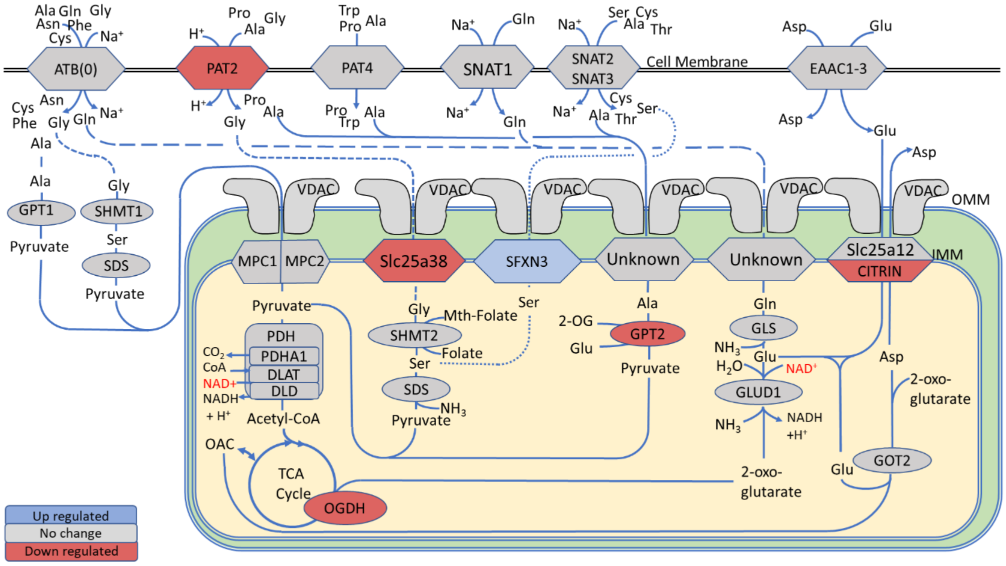

**Figure S8. Neutral amino acids pathways for alanine, glycine, serine, glutamine.** The schematics are partially based on the KEGG Pathway database.

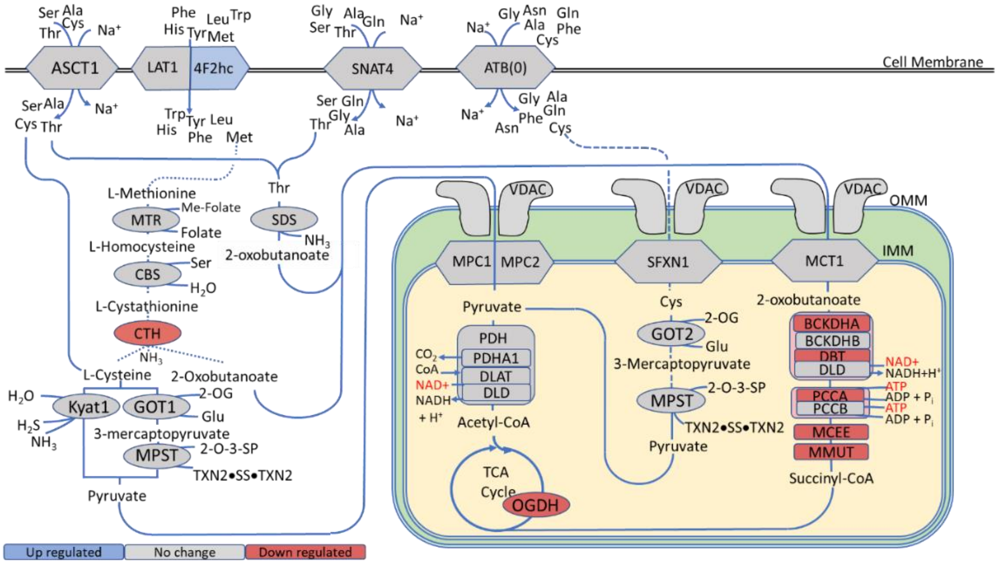

**Figure S9 Neutral amino acid pathways for cysteine, methionine and threonine affected by oxaliplatin.** The schematics are partially based on the KEGG Pathway database.

**Abbreviations to Figures S5-S7**. **Proteins abbreviations and encoded genes in parenthesis**: ACAA1A, 3-Ketoacyl-CoA thiolase (*Acaa1a*); ACAA2, mitochondrial 3-ketoacyl-CoA thiolase (*Acaa2*); ASCT1 and ATB(0), neutral amino acid transporters A and B(0) (*Slc1a4* and *Slc1a5*), AUH, mitochondrial methylglutaconyl-CoA hydratase (*Auh*); BCAT1/2, branched chain amino acid aminotransferases, (*Bcat1* and *Bcat2*); BCKDH, branched chain; BCKDHA and BCKDHB, branched chain α-keto-acid dehydrogenase subunits alpha and beta (*Bckdha* and *Bckdhb*); CBS, cystathionine beta-synthase (*Cbs*); CITRIN, electrogenic aspartate/glutamate antiporter SLC25A13, mitochondrial (*Slc25a13)*; CTH, cystathionine gamma-lyase (*Cth)*; DBT, mitochondrial lipoamide acyltransferase component of branched-chain alpha-keto acid dehydrogenase complex (Dbt); DLAT, dihydrolipoamide S-acetyltransferase pyruvate dehydrogenase complex component E2 *(Dlat*); DLD, mitochondrial dihydrolipoyl dehydrogenase (*Dld*); EAAC1 – EAAC3, excitatory amino acid transporters 1-3 (*Slc1a1*, *Slc1a2*, and *Slc1a3*); ECHS1, mitochondrial enoyl-CoA hydratase (*Echs1*); 4F2HC, 4F2 cell-surface antigen heavy chain (*Slc3a2*); GLS, glutaminase (*Gls*), GLUD1, mitochondrial glutamate dehydrogenase 1, (*Glud1*); GOT1, aspartate aminotransferase, cytoplasmic (*Got1*); GOT2, glutamic-oxaloacetic transaminase 2 (*Got2*); GPT1/2, alanine aminotransferases 1 and 2 (*Gpt1* and *Gpt2*); HADH, mitochondrial hydroxyacyl-coenzyme A dehydrogenase (*Hadh*); HADHA, mitochondrial trifunctional enzyme subunits alpha and subunit beta (*Hadha* and *Hadhb*); HIBCH, 3-hydroxyisobutyryl-CoA deacylase (hydrolase) (*Hibch*); HIBADH, 3-hydroxyisobutyrate dehydrogenase (*Hibadh*); HMGCL, mitochondrial hydroxymethylglutaryl-CoA lyase (*Hmgcl*); ICAD, isobutyryl-CoA dehydrogenase (*Acad8*); IVD, isovaleryl CoA dehydrogenase (*Ivd*); KYAT1, kynurenine—oxoglutarate transaminase 1 (*Kyat1*); LAT1, large neutral amino acids transporter small subunit 1 (*Slc7a5*); MCAD, mitochondrial medium-chain specific acyl-CoA dehydrogenase, (*Acadm*); MCCC1, mitochondrial methylcrotonoyl-CoA carboxylase subunits alpha and beta (*Mccc1* and *Mccc2*); MCEE, mitochondrial methylmalonyl-CoA epimerase (*Mcee*); MCT1, monocarboxylate transporter 1 (*Slc16a1*); MMUT, methylmalonyl-CoA mutase (*Mmut*); MMSDH, methylmalonic semialdehyde dehydrogenase (*Aldh6a1*); MPC1, mitochondrial pyruvate carrier 1 (*Mpc1*); MPC2, mitochondrial pyruvate carrier 2 (*Mpc2*); MPST, 3-mercaptopyruvate sulfurtransferase (*Mpst*); MTR, 5-methyltetrahydrofolate-homocysteine methyltransferase (*Mtr*); OGDH, 2-oxoglutarate dehydrogenase (*Ogdh*); PAT2, proton-coupled amino acid transporter 2 (*Slc36a2*); PAT4, neutral amino acid uniporter 4 (*Slc36a4*); PCCA and PCCB, mitochondrial propionyl-CoA carboxylase alpha and beta chains (*Pcca and Pccb*); PDH, pyruvate dehydrogenase E1 subunit alpha 1 (*Pdha1*); SBCAD, mitochondrial short/branched chain specific acyl-CoA dehydrogenase (*Acadsb*); SCAD, mitochondrial short-chain specific acyl-CoA dehydrogenase (*Acads*); SDS, L-serine dehydratase/L-threonine deaminase (*Sds*); SHMT1, serine hydroxymethyltransferase, cytoplasmic (*Shmt1*); SHMT2, mitochondrial serine hydroxymethyltransferase (*Shmt2*); SFXN1/3, sideroflexin-1 and sideroflexin-3 (*Sfxn1* and *Sfxn3*); SLC25A12, mitochondrial electrogenic aspartate/glutamate antiporter, (*Slc25a12*); Slc25a38, mitochondrial glycine transporter (*Slc25a38*); Slc25a44, solute carrier family 25 member 44 (*Slc25a44*); SNAT1-4, sodium-coupled neutral amino acid symporters 1-4 (*Slc38a1*, *Slc38a2*, *Slc38a3*, *Slc38a4*); Txn2, thioredoxin (*Txn2*); VDAC, mitochondrial outer membrane porin formed by voltage dependent anion channels 1-3 (*Vdac1*, *Vdac2*, and *Vdac3*). **Amino acid and metabolite abbreviations**: ADP, adenosine diphosphate; Ala, alanine; Asp, aspartate; ATP, adenosine triphosphate; Cys, cysteine; Folate, (6S)-5,6,7,8-tetrahydrofolate; Gln, glutamine; Glu, glutamate; Gly, glycine; Ile, isoleucine; Leu, leucine; 3-M-2OB, 3-methyl-2-oxobutanoate; 3-M-2OP, (S)-3-methyl-2-oxopentanoate; 4-M-2OP, 4-methyl-2-oxopentanoate; Me-Folate, (6S)-5-methyl-5,6,7,8-tetrahydrofolate; Mth-Folate, (6R)-5,10-methylene-5,6,7,8-tetrahydrofolate; NAD+, nicotinamide adenine dinucleotide (oxidized); NADH +, nicotinamide adenine dinucleotide (reduced); OAC, oxaloacetic acid; 2-OG, 2-oxoglutarate (alpha-ketoglutarate); Phe, phenylalanine; Pi, inorganic monophosphate; Pro, proline; Ser, serine; Thr, threonine; Trp, tryptophan; Val, valine. **Other abbreviations**: TCA cycle, tricarboxylic acid cycle (citric acid cycle); OMM, outer mitochondrial membrane; IMM, inner mitochondrial membrane. The pathways are derived from the KEGG database. Blue – upregulated genes, red downregulated genes, grey – not affected genes.

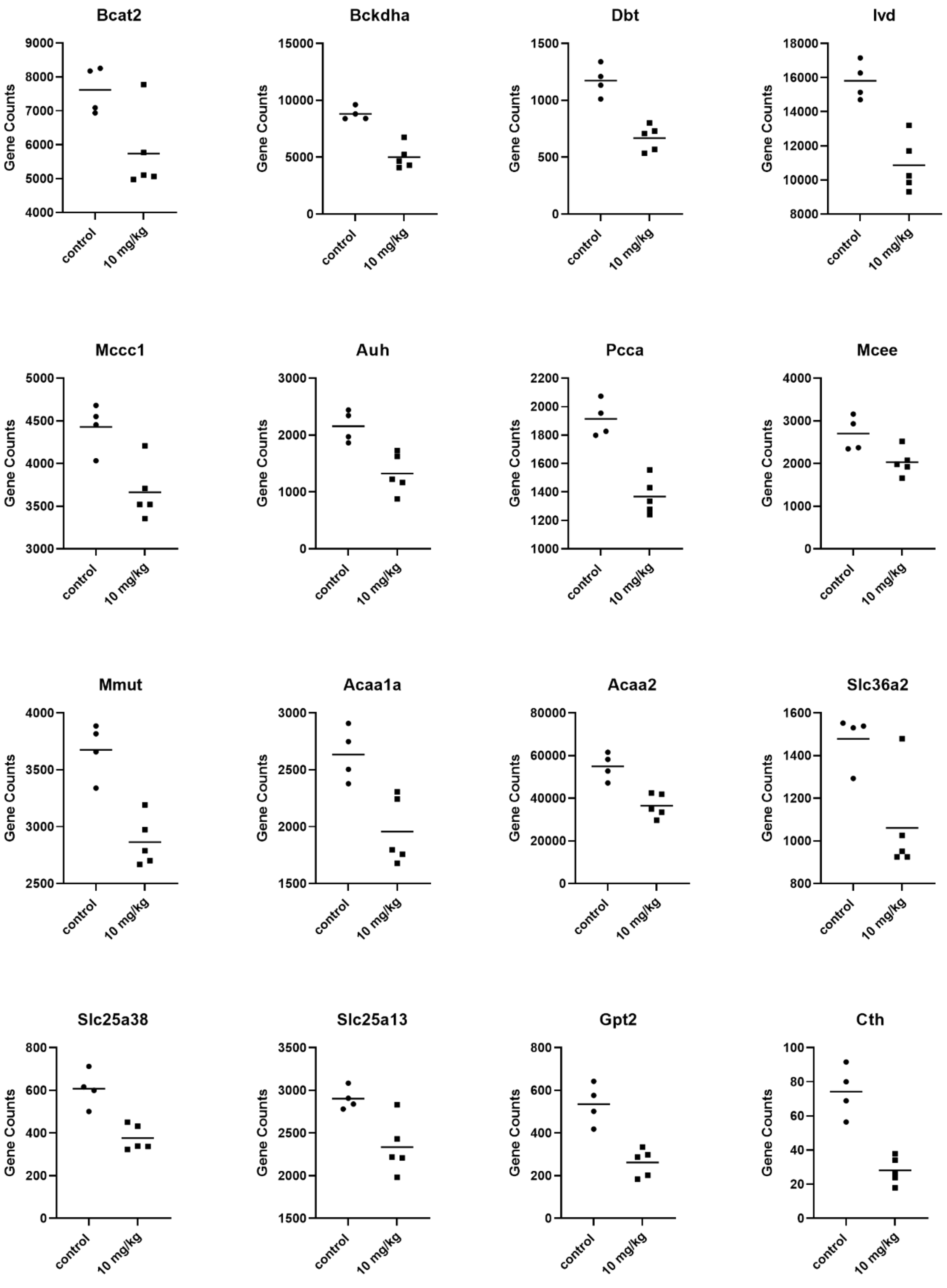

**Figure S10** Oxaliplatin downregulates genes involved in the catabolism of amino acids: *Bcat2* (FC = -1.33, FDR = 0.04), *Bckdha* (FC = -1.76, FDR = 4.10E-06), *Dbt* (FC = -1.76, FDR =

5.16E-06), *Ivd* (FC = –1.46, FDR = 2.9E-4), *Mccc1* (FC = –1.21, FDR = 0.04); *Auh* (FC = –1.63, FDR = 3.02E-3), *Pcca* (FC = –1.40, FDR = 1.49E-4), *Mcee* (FC = –1.33, FDR = 0.04), *Mmut* (FC = –1.28, FDR = 3.13E-3), Acaa1a (FC = −1.35, FDR = 0.01) *Acaa2* (FC = −1.51, FDR = 3.69E-4), *Slc36a2* (FC = -1.39, FDR = 0.02), *Slc25a38* (FC = -1.61, FDR = 3.57E-04), *Slc25a13* (FC = -1.24, FDR = 0.05), *Gpt2* (FC = −2.05, FDR = 2.58E-5), *Cth* (FC = −2.63, 1.50E-04).

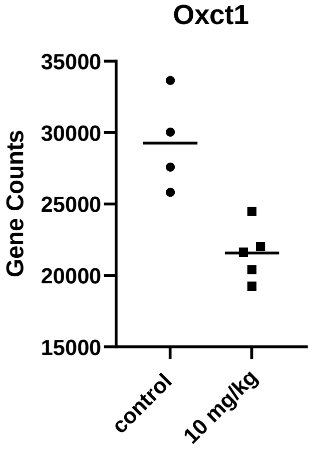

**Figure S11** Oxaliplatin downregulates ketone body metabolism by downregulating *Oxct1* (FC = −1.3, FDR = 1.72E-3)

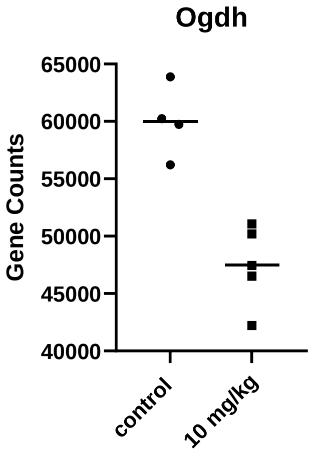

**Figure S12** *Ogdh* exhibited statistically significant down-regulation (FC =-1.28, FDR = 2.88E-3)

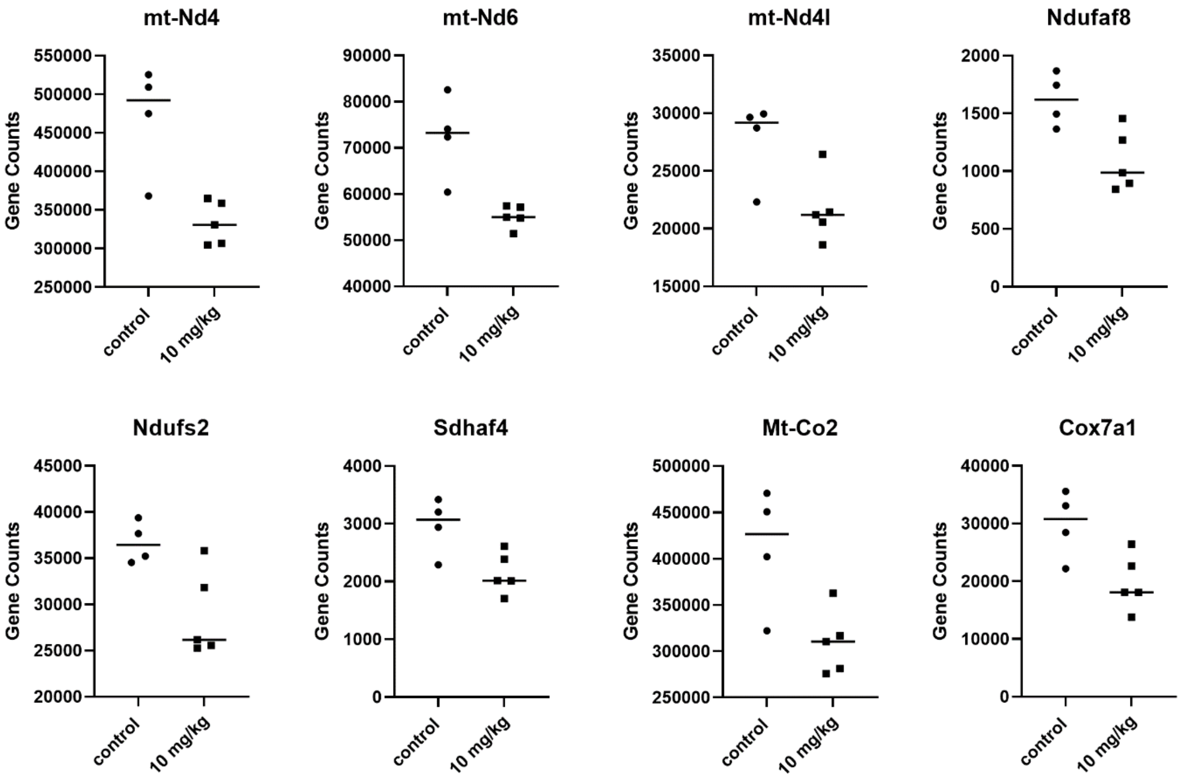

**Figure S13** Genes that provide instructions for Electron Transport Chain (ETC) are downregulated in oxaliplatin treated mice. Complex 1: *mt-Nd4* (FC = −1.41, FDR = 1.43E-3); *mt-Nd6* (FC = -1.31, FDR = 3.25E-3); *mt-Nd4l* (FC = −1.28, FDR = 0.05); *Ndufaf8* (FC =−1.49, FDR = 0.01), *Ndufs2* (FC = −1.27, FDR = 0.05); Complex II: *Sdhaf4* (FC =−1.38, FDR = 0.02); Complex IV: *Mt-Co2* (FC = -1.33, FDR = 0.02) and *Cox7a1* (FC = −1.51, FDR = 0.02). Uncoupling protein gene *Ucp2* is overexpressed: (FC = 2.46, FDR = 4.54E-6)

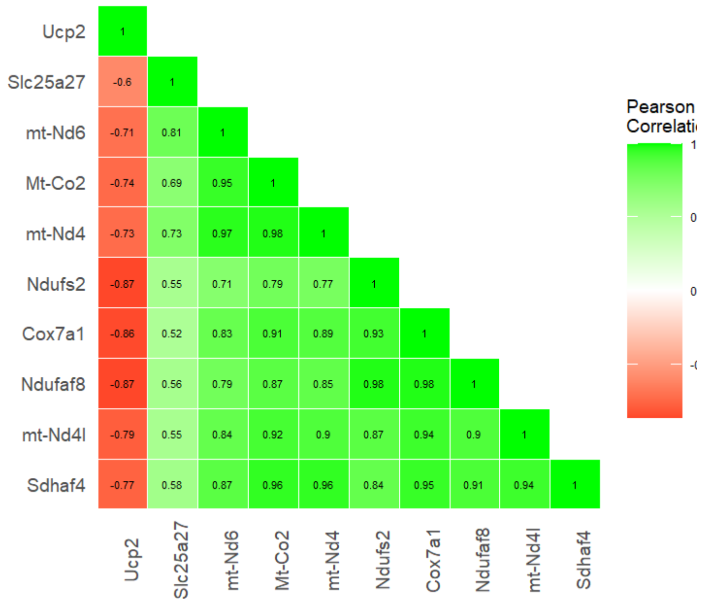

**Figure S14** Oxidative phosphorylation (Electron Transport Chain): correlation matrix of differentially expressed genes

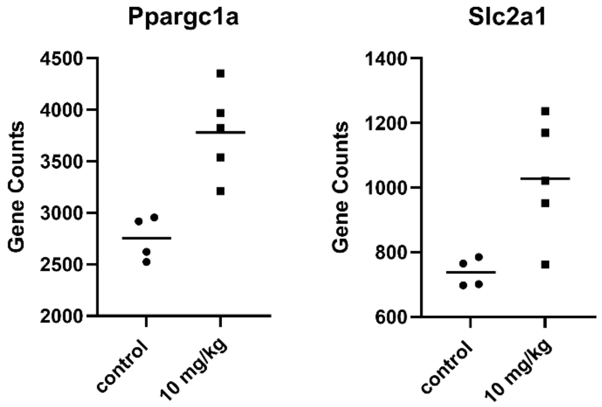

**Figure S15** Oxaliplatin upregulates the energy-sensing enzyme *Ppargc1a* (FC=1.37-fold increase, FDR = 1.39E-3). Increase of glucose transport by *Slc2a1* (GLUT1) (1.39-fold increase, FDR = 0.01) suggests a partial switch to glycolysis.

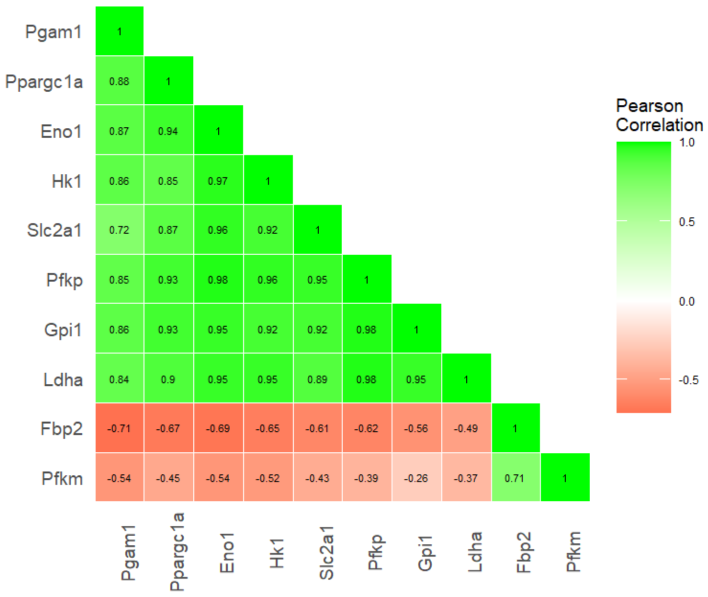

**Figure S16** Glycolysis: correlation matrix of differentially expressed genes

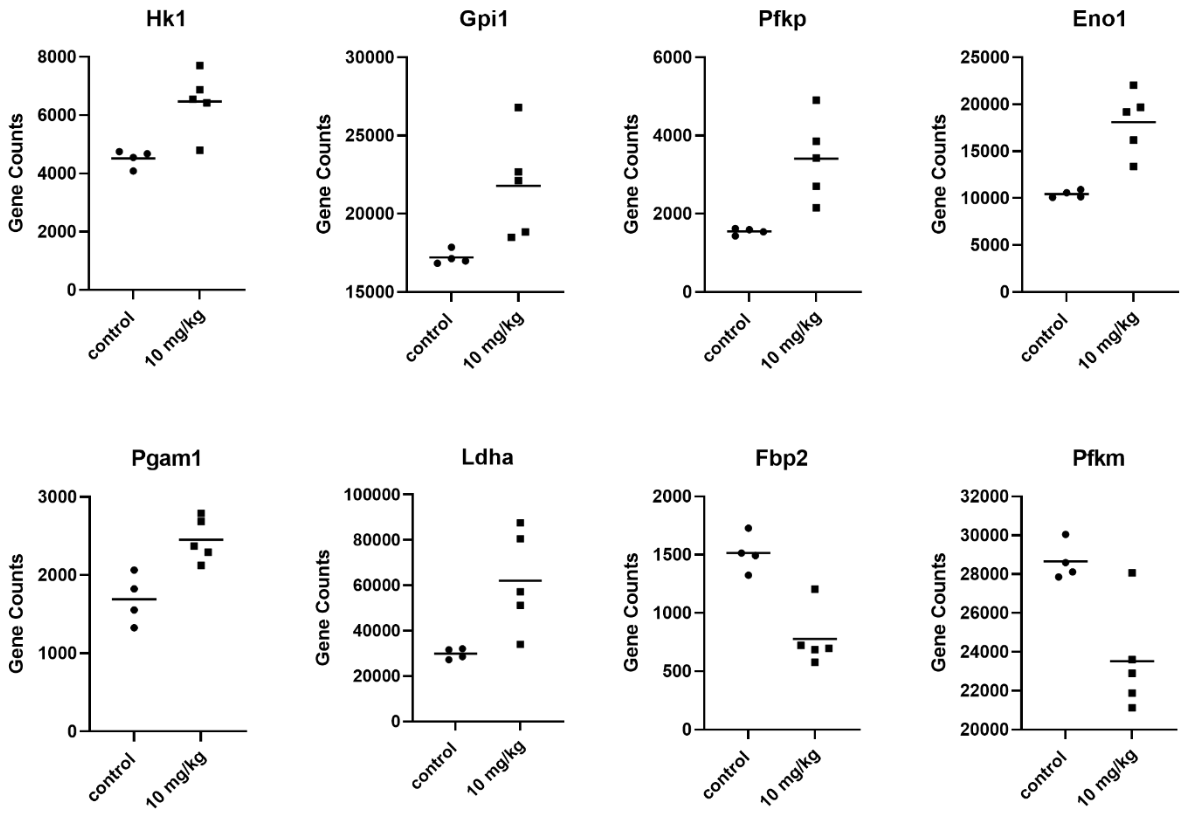

**Figure S17** Oxaliplatin leads to significant changes in gene expression in glycolysis pathway. Upregulated: *Hk1* (FC = 1.43, FDR = 2.29E-3), *Gpi1* (FC = 1.27, FDR = 0.04, *Pfkp* (FC = 2.2, FDR = 1.2E-6), *Pgk1*, FC = 1.27, FDR = 0.04), *Eno1* (FC = 1.74, FDR = 1.84E-6), *Pgam1* (FC =

1.45, FDR = 3.97E-3), *Ldha* (FC = 2.08, FDR = 7.89E-05); *Fbp2* (FC = −1.95, FDR = 4.18e-5) and *Pfkm* (FC = −1.22, FDR = 0.04) are downregulated.

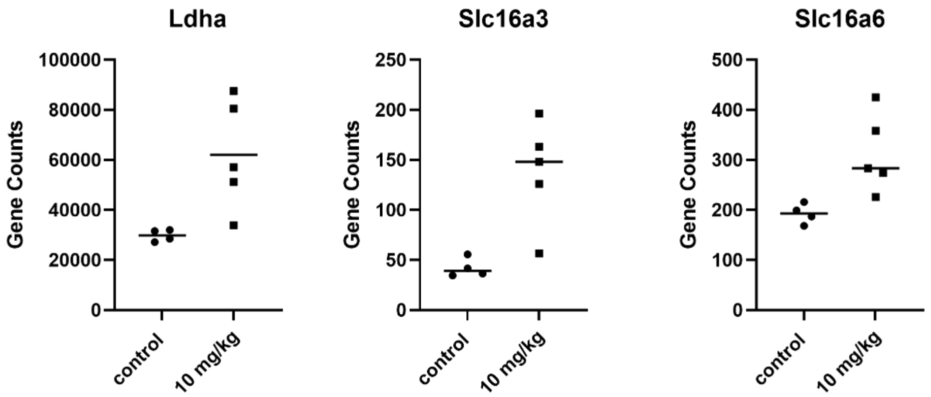

**Figure S18** Oxaliplatin upregulates genes related to lactate conversion in the Cori cycle: *Ldha* (FC = 2.08, FDR = 7.89e-5), *Slc16a3* (FC = 3.27, FDR = 1.63e-5); *Slc16a6* (FC = 1.62, FDR = 5.84E-3)

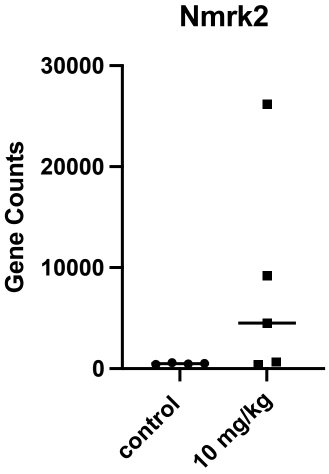

**Figure S19** Oxaliplatin treated mice show robust increase in the expression of *Nmrk2* (FC = 16.44, FDR = 1.4E-3) and other enzymes from the NAD pathway

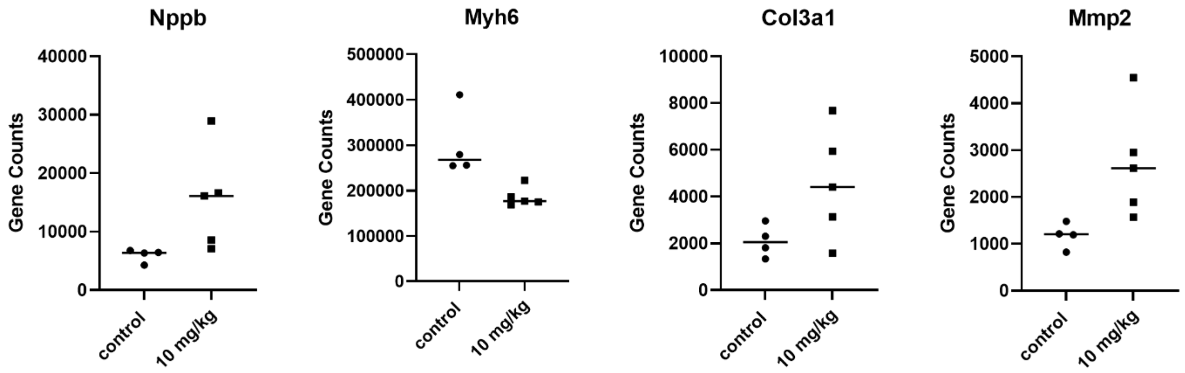

**Figure S20** Oxaliplatin induces changes in the genes encoding markers of the heart damage. *Nppb* (FC = 2.59, FDR = 0.01), *Myh6* (FC = −1.61, FDR = 3.275e–4), *Col3a1* (FC = 2.17, FDR = 0.07) and *Mmp2* (FC = 2.30, FDR = 1.61e-4)
